# timePlex: Increasing mass spectrometry throughput using time-encoded sample multiplexing

**DOI:** 10.1101/2025.05.22.655515

**Authors:** Jason Derks, Kevin McDonnell, Harrison Specht, Daniel J. Geiszler, Michael P. Agius, Xiaofeng Xie, Jarrod J. Sandow, Liang Chen, Lina Kozhaya Warinner, Nathan Wamsley, Sarah N. Sipe, Peyton Stewart, Maddy Yeh, Nikolai Slavov

**Author notes:** Correspondence &. ∈Data, code & protocols: parallelsq.org/timePlex.

## Abstract

Liquid chromatography-mass spectrometry (LC-MS) can enable precise and accurate quantification of analytes at high-sensitivity, but the rate at which samples can be analyzed remains limited. Throughput can be increased by data-independent-acquisition (DIA) using short separation times and by multiplexing samples in the mass domain with plexDIA; yet multiplexing along one dimension scales throughput only linearly with plex. To enable combinatorial scaling of proteomics throughput, we developed a complementary DIA strategy that multiplexes samples in the time domain, termed ‘timePlex’. timePlex staggers and overlaps the separation periods of individual samples. This strategy is orthogonal to isotopic multiplexing, which enables combinatorial multiplexing in mass and time domains, and thus multiplicatively increased throughput. We demonstrate this with a 27-plex (3-timePlex and 9-plexDIA) exceeding 1,200 samples (4M protein datapoints) per day. These results establish timePlex as a framework for multiplexing both label-free and labeled samples that supports sensitive and accurate protein quantification and combinatorial scaling.

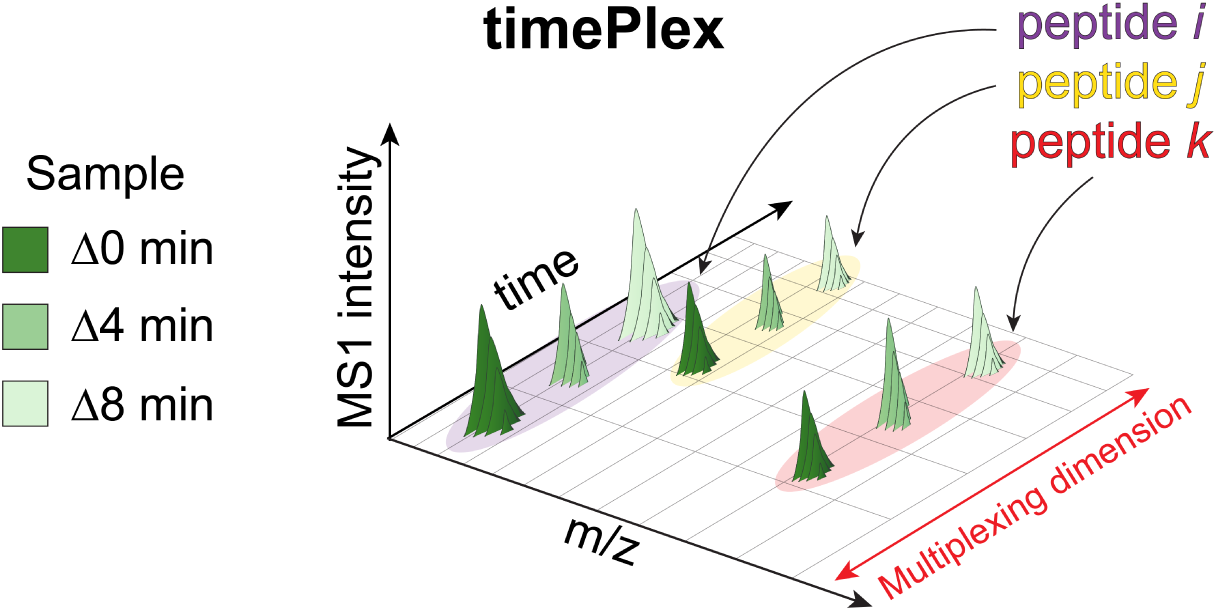

## Introduction

Liquid chromatography-mass spectrometry (LC-MS) proteomics methods can achieve accurate quantification while analyzing over 50 samples per day^1–3^. This has been made possible due to advances in liquid chromatography that have enabled robust and repeatable separations of chemical analytes from complex mixtures^4–7^, mass spectrometry instrumentation and data acquisition^8–13^, and data interpretation approaches^14–17^. Together, these advances have positioned LC-MS proteomics as a powerful tool in quantitative biology. However, the scale at which such datasets can be generated remains constrained by time and costs associated with LC-MS data acquisition. As such, there is need for increased throughput of data acquisition to generate datasets that will empower biological investigations.

Strategies to increase the throughput of LC-MS proteomics generally aim to 1) increase the number of LC-MS runs completed per unit time^2,3,18,19^, or 2) increase the number of samples acquired per LC-MS run as achieved through sample multiplexing^20–22^. Both strategies are complementary approaches to increase LC-MS throughput^22–25^. The former has inherent limitations, in that the number of quantifiable points per peak becomes limiting as LC-MS separations are shortened; this challenges quantitative accuracy even with the fastest instruments and duty cycles^26^. This challenge can be addressed by sample-multiplexing that can enable high throughput while using separation times long enough to support multiple points per peak and thus quantitative accuracy.

Sample multiplexing is commonly achieved through chemically barcoding samples with isotopic mass offsets to be pooled and acquired in parallel within the same run. The samples are separable at MS1 level with isotopologous mass offsets such as mTRAQ, PSMtag, or SILAC among others^27–32^, or at the MS2- and MS3-level with isobaric mass-tags such as TMT and iTRAQ^33,34^. While 35-plex TMTpro currently offers the highest multiplexing capacity^35,36^, quantifiable proteomic coverage per unit time is fundamentally limited as ‘reporter ions’ cannot be used to quantify precursors of chimeric spectra. Thus, quantification based on reporter ions is limited to quantifying one precursor per MS2 scan. Alternative approaches such as data-independent acquisition (DIA)^37,38^ parallelize precursor analysis through the intentional generation of chimeric spectra, enabling greater depth of coverage per unit time; such approaches are amenable to non-isobaric isotopologous multiplexing^39^. Indeed, multiplexed DIA (plexDIA) has been shown to multiplicatively increase throughput at high depth of coverage and accuracy^20,21,40^.

Sample multiplexing can be further increased based on an orthogonal and relatively unexplored dimension for multiplexing proteomics samples: time. Samples can undergo parallelized and independent separations which are offset in time, encoding sample-specificity in the time-domain. Temporal encoding has been implemented for sample multiplexing based on multi-segment capillary electrophoresis (CE) and targeted analysis of relatively few analytes^41–43^. Here, we explore the potential of using time-domain multiplexing for parallelized data acquisition using DIA. This approach requires new strategies to interpret the resulting multiplexed mass spectra. Such an approach may scale not only sample throughput defined as samples per day, but also throughput defined as quantified proteins per unit time.

Here we demonstrate two implementations of timePlex that enable the acquisition of three samples in parallel (3-timePlex). To interpret the unique LC-MS data generated by timePlex, we developed a software module in JMod^44^. Furthermore, we applied combinatorial multiplexing of 3-plexDIA and 3-timePlex to acquire 9 samples in parallel per LC-MS run, and 9-plexDIA^32,44^ and 3-timePlex for 27-plex analysis. The two implementations establish a framework for multiplexing samples in time that we expect will expand to higher plex, generalize to other implementations, and enable a combinatorial increase in proteomics throughput when paired with isotopic labeling approaches such as plexDIA.

## Results

### Multiplexing in the time and mass domains

The timePlex concept and its relationship to other LC-MS approaches is illustrated in Fig. 1a with elution curves for three precursors. Acquiring data by non-multiplexed DIA limits the rate of data acquisition to a single sample per run. Multiplexing in the mass dimension through mass-encoded offsets, as performed in plexDIA, enables a linear increase in the rate of sample acquisition with plex. Here we introduce a complementary approach, timePlex, by multiplexing samples using time-encoded offsets to likewise increase sample-throughput. Due to their orthogonality, timePlex and plexDIA can be used in combination to increase proteomics throughput multiplicatively.

**Figure 1.**
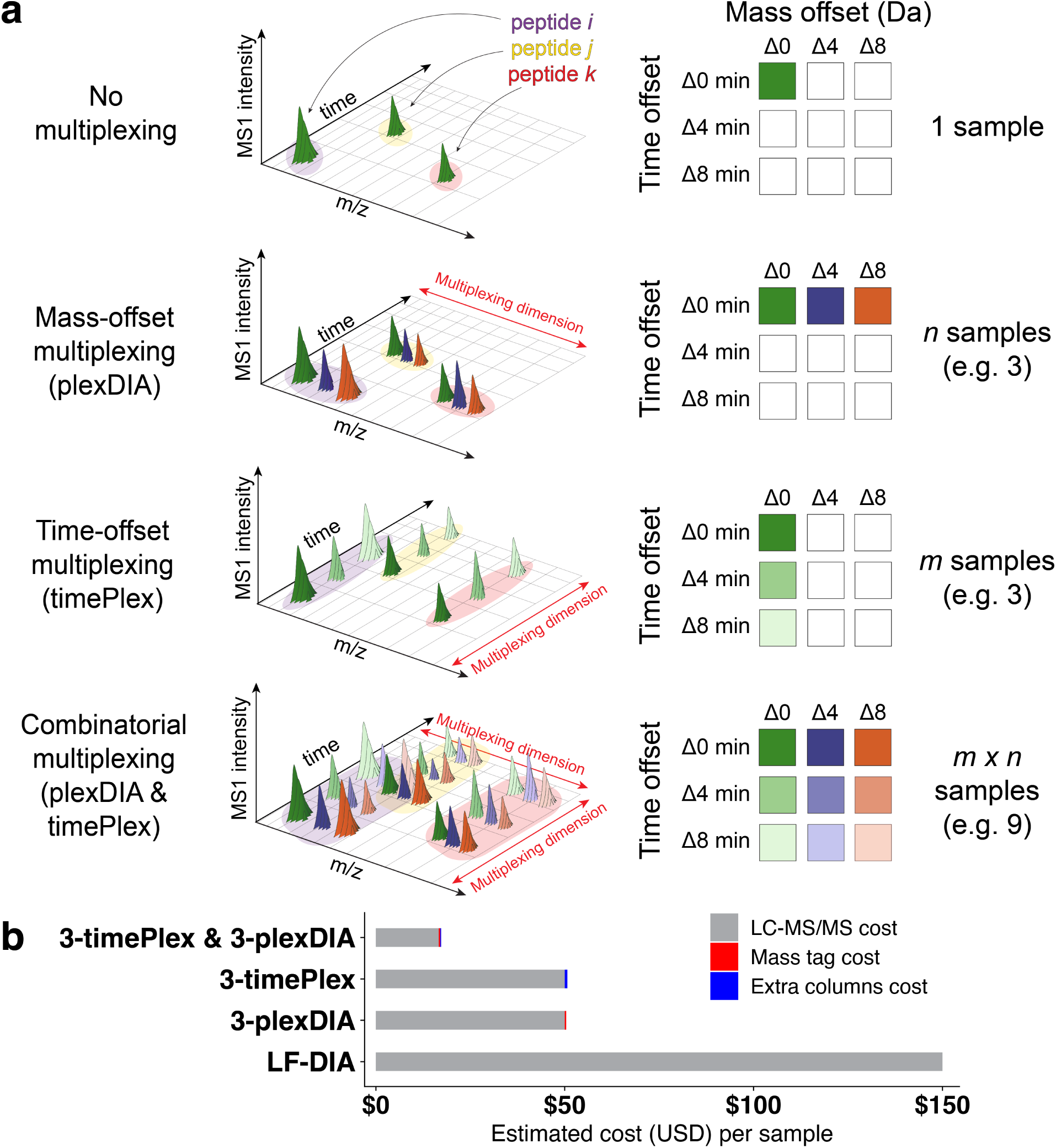
Multiplexing in the time and mass domains is combinatorial. **a** Illustration of precursor elution curves for no multiplexing, plexDIA, timePlex, and combined timePlex & plexDIA data acquisitions. Combinatorial timePlex & plexDIA enables multiplicative scaling of the number of samples acquired / run. **b** Estimated cost for LC-MS analysis per sample for different levels of multiplexing not including time spent on sample-loading overheads. The total cost per sample is estimated assuming $150/hr for LC-MS/MS, approximately $1k/column with an average 1,000-run lifespan, and the cost to label 1 *µ*g of protein with a commercially available tag (approximately $0.36 per sample).

Multiplexing nearly proportionally reduces the cost of data acquisition because the expense of LC-MS instrumentation and operation exceeds that of mass tags and LC columns. timePlex and plexDIA may be used in combination, increasing throughput and cost-saving multiplicatively with plex, as shown in Fig. 1b. Here, we seek to develop an approach for timePlex proteomics and benchmark its combinatorial potential when paired with plexDIA.

### Implementing timePlex proteomics data acquisition

Multiplexing through temporal encoding can be implemented through several distinct approaches. Here, we aim for a proof of principle with two accessible implementations with different column types illustrated in Fig. 2a and expect significant improvements in future implementations. In both approaches, the gradient flow originating from a single LC-system is split to separate columns. Time offsets are encoded by adjusting the volume of each sample’s respective capillary transfer line. In the first implementation, termed ‘single-emitter’, the flow from each column is rejoined to a single emitter for electrospray ionization (ESI). In the second approach, termed ‘separate-emitters’, the flow remains separate and they undergo separate ESI from their respective emitters, the setup for which is shown in Supplementary Fig. 1. In both cases, the resulting MS data will include multiple observations of the same peptides, separated by systematic time offsets which are encoded by the different travel paths, Fig. 2b.

**Figure 2.**
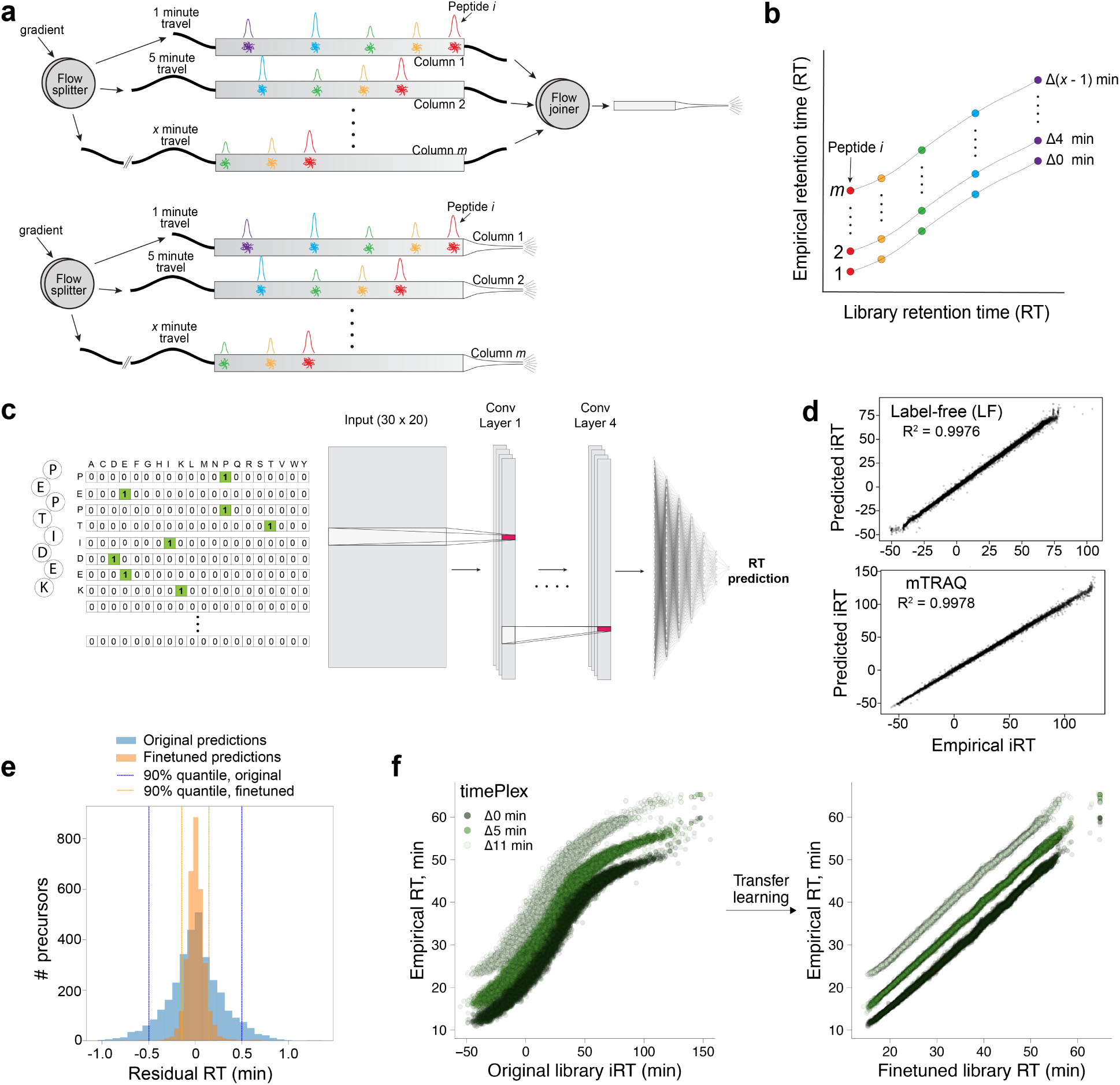
Enabling timePlex analysis through accurate RT prediction. **a** Design of two timePlex implementations with a single emitter (top) and separate emitters (bottom). The time-offsets are encoded through differences in the volumes of the transfer lines. **b** Conceptual illustration of repeated observations of peptide species systematically offset in time. **c** Illustration of our RT predicting model. Peptide sequences were one-hot encoded and a hybrid convolutional neural network and deep neural network architecture was trained to predict retention times for label-free (LF) and mTRAQ-labeled peptides. Models were trained on 91,657 LF peptides or 56,645 mTRAQ-labeled peptides. **d** Test data, which were withheld from training, were used to evaluate the performance of the RT predictor. **e** To evaluate the model’s ability to fine-tune RT’s from new data, residuals were computed between the original RT predictions of the model (shown in blue), and after finetuning on approximately 3,200 peptides (shown in orange); the plotted data is the test-set withheld from fine-tuning. The 90th percentile boundaries are shown with orange and blue lines. **f** The original library iRTs predicted during DIA-NN *in-silico* library generation of LF data were used to visualize a typical starting point of RT accuracies, as shown in the left panel. For each run, our model uses the initial search results to fine-tune RTs to be used in the final search. The final search output is shown in the right panel with the library RTs for the middle timePlex channel.

In these implementations, samples are sequentially loaded and travel through the same sample loop and parts of the same transfer lines to reach their designated columns during sample-loading. Schematics for implementing a 3-timePlex are shown in Supplementary Fig. 2. During sample loading, peptides may bind nonspecifically to surfaces in the common flowpath, leading to sample-carryover into unintended columns. To mitigate or prevent this altogether, samples were resuspended in 0.015% DDM prior to sample-loading. Carryover was undetectable for the vast majority of peptides; when detected, it was limited to a median of *<*3% of the intensity of the intended time channel, Extended Data Fig. 1.

### Interpreting timePlex proteomics data

Interpretation of timePlex data depends on highly predictable elution times of analytes (e.g., peptides). While retention time (RT) offsets are less precise and reproducible than *m/z* offsets, they can afford reliable sample assignment if they are engineered to be large enough to account for the accuracy of RT prediction. To analyze time-encoded multiplexed data, we developed a specialized timePlex module in JMod^44^. The module implements time channel detection followed by a time channel-dependent main search. The time channel detection determines the times when precursors from each channel are expected. Then, these times are used to perform the main search. The time channel detection starts by identifying peptide sequences that are detected across all timePlex channels; this allows for assigning peptides to a given timePlex channel based on the temporal order of each peptide’s multiple elutions, each corresponding to a timePlex channel. These preliminary identifications are used to finetune the retention-times of the initial library, which are then aligned to each time channel separately. This updated library is then used for the main search where peptides can be confidently assigned to a timePlex channel. This approach remains flexible to differences in time-offsets between multiple time channels, as the alignment is flexible to fitting each timePlex channel. As a result, the user does not need to specify the time spacing between samples, only the number of timePlex channels.

While precursors across time channels are systematically separated in time, accurately assigning each identified precursor to a sample becomes challenging when a given peptide is not observed in all time channels. This difficulty stems from inaccuracies in library retention times. To address this, we built a retention-time predictor^45–52^ to fine-tune the library retention times on a per run basis^53^. The model architecture implemented in this work is a convolutional neural network ending in fully connected layers of a feedforward deep neural network as illustrated in Fig. 2c. A summary of the model architecture is shown in Supplementary Fig. 3. LF and mTRAQ models were trained on fewer than 100k peptides each; test-data was withheld to evaluate the accuracy of the models for unseen mTRAQ and LF peptides. We observed strong agreement between predicted and empirical iRTs (*R*^2^ *>* 0.997) for both label-free and mTRAQ-labeled peptides, as shown in Fig. 2d, which suggests these models accurately predict retention times, and may be suitable for transfer learning.

Data generated outside of these training and test sets (different gradient, instruments, and column batch) was used to evaluate the potential for the models to refine library retention times given new data acquired under different conditions. We found that the original model was able to predict 90% of the data within 30.0 seconds of the empirical RT, and after fine-tuning it improved to within 8.8 seconds as shown in Fig. 2e. The improved RT accuracy stands to benefit DIA search performance due to increased specificity of retention time search spaces, even in non-timePlex applications, as previously shown by Wallmann *et al.*^53^

Fine-tuning retention times on a per run basis is of even greater importance with timePlex data where inaccurate library retention times may lead to ambiguous time channel assignment of peptides. To evaluate fine-tuning in real timePlex applications, we compared the iRTs generated from DIA-NN *in-silico* library prediction to the empirical RTs in a LF 3-timePlex experiment, Fig. 2f. Despite the consistently encoded retention time offset, the 3-timePlex data display overlap between timePlex channels. This ambiguity is due to inaccuracies in the library RTs, which we resolve through updating library RTs based on initial search results using transfer learning. Using the distribution of residuals between predicted and empirical RTs of this initial search, thresholds are dynamically set to confine the RT-space where a precursor will be considered, shown in Supplementary Fig. 4. This fine-tuning step enhances the ability of properly assigning sample-origins to each precursor. The resulting fine-tuned and searched data produce clear temporal separation between the timePlex channels, as shown in the right panel of Fig. 2f.

### Benchmarking timePlex proteomic coverage through mixed-species standards

To investigate the potential for false identifications in 3-timePlex, we acquired data on a Thermo Orbitrap Exploris 240 from samples containing *H. sapiens* (human) protein digests present at 10 ng in all three time channels and *S. cerevisiae* (yeast) digests only present at 5 ng in the last two. These data were searched with JMod using a *H. sapiens*, *S. cerevisiae*, and *A. thaliana* spectral library, searching for peptides from all species in each sample. Across all channels, this workflow identified over 25k human precursors and an average of 40 *A. thaliana* precursors per channel, consistent with the computationally set FDR of 1%, Fig. 3a. It also identified thousands of yeast precursors in the time channels containing yeast digest but only 371 lowly abundant yeast precursors in the time channel missing yeast. To more directly display the raw data supporting these results, we visualized the extracted ion chromatograms for a precursor corresponding to human GAPDH and its yeast homolog TDH1, Fig. 3b. As expected, signal was present for each human precursor across time channels, while the yeast precursor was present at an intensity of *<*1% in the Δ0 min time channel relative to the Δ4 and Δ8 min time channels, consistent with column carryover rather than timePlex misalignment.

**Figure 3.**
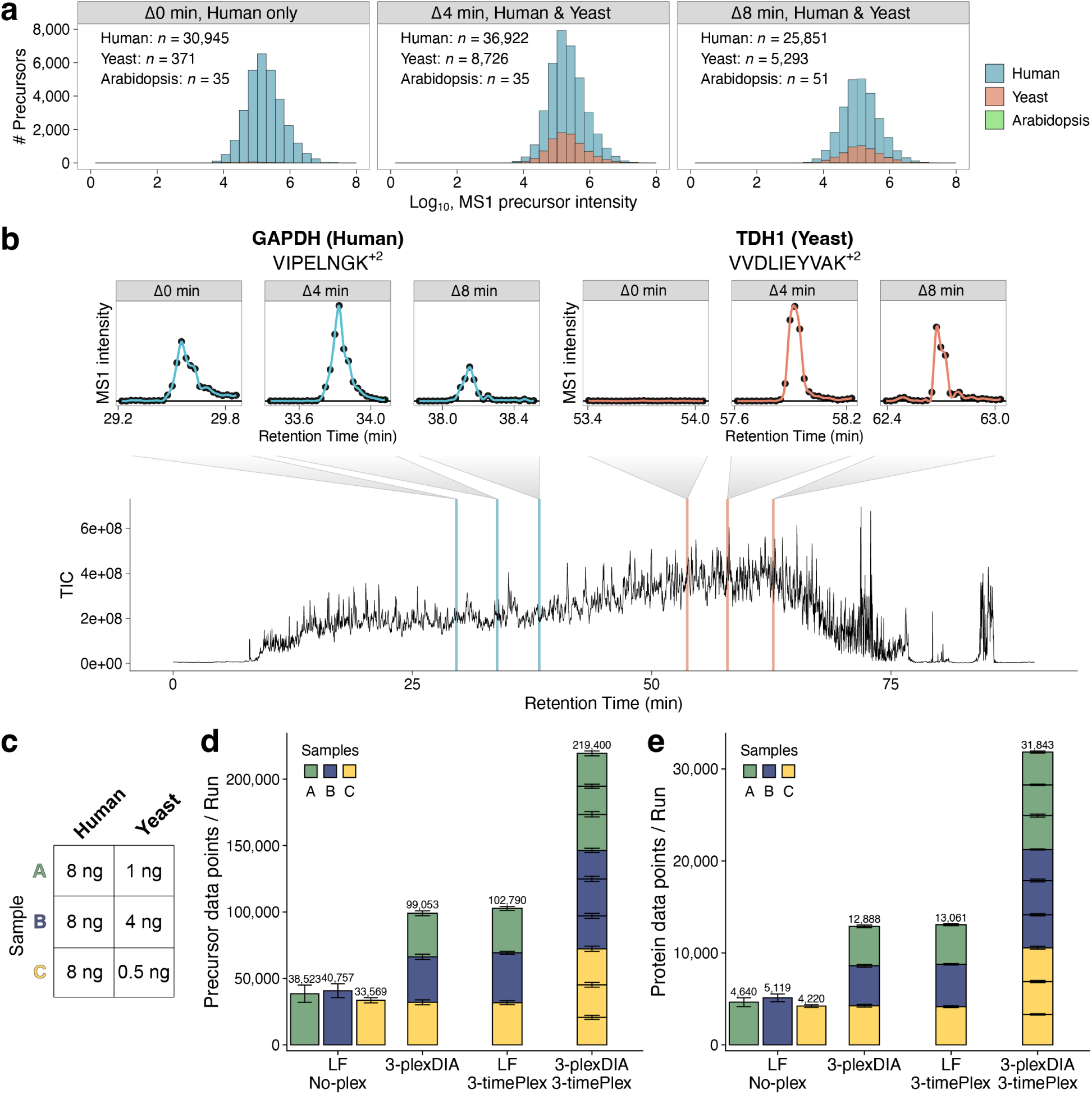
Benchmarking timePlex protein coverage and throughput scalability. **a** LF 3-timePlex data was acquired on an Orbitrap Exploris 240 from three time channels (Δ0 min, Δ4 min, and Δ8) containing human only, human & yeast, and human & yeast, respectively. The data was searched with a human, yeast, and Arabidopsis spectral library. The number of precursor identifications is shown in each sub-panel for each species in addition to overlapping histograms of MS1 intensities for all precursors identified. **b** The total ion chromatogram (TIC) of the missing-species run is displayed in addition to extracted ion chromatograms for a human GAPDH precursor and a yeast homolog of GAPDH (TDH1). The data show signal for human precursors across all three time channels and yeast signal at *<*1% intensity in the Δ0 min time channel. **c** Sample composition for benchmarking protein coverage and quantitative accuracy. **d-e** Precursor data points **d** and protein data points **e** identified per run for LF no-plex, 3-plexDIA, LF 3-timePlex, and 3-timePlex & 3-plexDIA with the ‘separate emitters’ approach; error bars reflect standard error.

Next, we sought to benchmark the depth of coverage achieved with 3-timePlex and combinatorial 3×3 plexDIA & timePlex, relative to LF “no-plex”. To accomplish this, we created mixed-species standards of human and yeast proteomic digests, in which human composition was held constant and yeast was variably spiked at known differential abundances as standard practice to benchmark quantitative accuracy^54^. The sample compositions are shown in Fig. 3c. These samples were analyzed using an Orbitrap Exploris 240. Before comparing timePlex to established methods, we acquired non-timePlex data with and without the timePlex setup to directly evaluate the impact of the ‘separate emitters’ timePlex implementation on ion intensities and proteome coverage. This experiment was set up to evaluate the effect of increasing ESI plume density in timePlex where the flow rate for the 1 column approach was 200 nL/min and 3 times higher (600 nL/min total) for the 3 columns setup. Results in Extended Data Fig. 2a-c show that the 3-column timePlex ‘separate emitters’ approach results in an ion intensity loss of approximately 3-fold and the identification of approximately 20% fewer precursors. However, we found that lowering the flow rate to 66 nL/min/column produced equivalent intensities and coverage for the 3-column timePlex setup and the standard single column setup. To avoid the influence of other factors and more clearly evaluate the timePlex data type, the benchmarking experiments were performed with identical LC-MS setups regardless of the methodology used for acquisition.

To control for the influence of the timePlex implementation described above, we evaluated the proteomic depth of coverage for plexDIA, timePlex, and combinatorial plexDIA & timePlex relative to label-free (LF) no-plex data using the timePlex setup. The implementation with separate emitters was acquired with 8.5-12 ng human and yeast peptide samples on a Thermo Orbitrap Exploris 240. LF no-plex quantified an average of 37,616 precursors and 4,660 proteins per run between samples A, B, and C, Fig. 3d,e. 3-plexDIA quantified 99,053 precursor data points/run and 12,888 protein data points/run and 3-timePlex quantified 102,790 precursor data points/run and 13,061 protein data-points per run. This corresponds to an average of 33,018 and 34,263 precursors/sample, and 4,296 and 4,354 proteins/sample for 3-plexDIA and 3-timePlex respectively. Therefore, both multiplexing approaches quantified approximately 2.6 - 2.7-fold more precursor data points/run and 2.8-fold more protein data-points/run than LF no-plex. The combination of 3×3 plexDIA & timePlex identified 219,400 precursor data points/run and 31,843 protein data-points/run for an average of 24,378 precursors/sample and 3,538 proteins/sample. Thus, 3×3 plexDIA & timePlex with the current implementation of multi-emitters resulted in 5.8-fold more precursor data points/run and 6.8-fold more protein data-points per run compared to LF no-plex.

We observed similar proportional scaling of coverage for the single emitter merged flow approach with *µ*PAC columns, Extended Data Fig. 3a-b. With this setup, LF no-plex identified an average of 29,771 precursors and 3,769 proteins between samples A, B, and C per run. 3-timePlex identified an average of 60,884 precursor data points per run and 9,009 protein data points per run, producing approximately 2.0-fold and 2.4-fold as many precursor and protein data points per run compared to LF no-plex. The combination of 3-timePlex & 3-plexDIA identified a total of 163,012 precursor data points per run and 23,749 protein data points per run on average, thus producing approximately a 5.5-fold and 6.3-fold increase in the number of precursor and protein data points identified per run, respectively, compared to LF no-plex. This scaling is likely to be higher with faster instruments that can better meet the increased ion complexity of the multiplexed samples.

Search softwares that jointly model fragments, such as JMod and Specter, rely on accurate spectral libraries^44,55^. As the spectral complexity of DIA data increases, there may be increased sensitivity to spectral library quality. To investigate this potential, we searched 3-plexDIA and 3×3 plexDIA & timePlex data with two libraries, one optimized for mTRAQ-labeled peptides and another optimized for label-free peptides. Because the label-free fragmentation patterns do not accurately reflect mTRAQ-labeled peptide fragmentation, we identified 28.1% more precursors with the mTRAQ-optimized library than when using the label-free library, Extended Data Fig. 4a-b. Similar search of the higher plex 3×3 plexDIA & timePlex data with a mTRAQ-optimized library identified an even greater proportion of precursors on average, with a 37.2% increase. The larger increase for the higher plex data suggests that spectral library quality may become increasingly important at increasingly high plex; this may partially explain the proportionally decreased coverage per sample in 3×3 plexDIA & timePlex, which may be mitigated through development of spectral libraries which more closely reflect empirical spectral fragmentation patterns.

### Investigating and correcting chromatographic biases

We evaluated column-specific differences in elution profiles, shown in Extended Data Fig. 5a, as differences have the potential to impact timePlex coverage per time channel. The columns within each implementation performed comparably, aside from one column from the ‘single emitter’ implementation which produced 2-fold broader elution peaks. As a result, the number of precursor identifications was similar within each implementation, except for the third column with the single-emitter approach, Extended Data Fig. 5b-c.

Additionally, we observed RT-dependent differences in relative precursor ion intensities for timePlex data. As shown in Extended Data Fig. 6a-b, discrepancies between LF timePlex and LF no-plex quantitation are primarily correlated with timePlex total ion chromatogram (TIC) intensities, suggesting the discordance is best explained by timePlex induced variability. Additionally, TIC and precursor ratios of timePlex and LF no-plex data covary, Extended Data Fig. 6c, suggesting the timePlex induced effects are variable with RT. Fortunately, these differences are systematic and thus likely correctable. We therefore perform a quantitative correction to stabilize precursor quantities across the gradient, whereby MS1 and MS2 intensities in each time channel are corrected by a LOESS regression of log-transformed intensities against retention time. Time-channel specific variability is further reduced through a second LOESS regression on relative precursor deviations between time channels across the gradient. MS1- and MS2-level ratios between timePlex and LF no-plex are shown in Extended Data Fig. 6d-g before and after correction, and indicate approximately a 7-10% decrease in standard deviation and more identical quantitation across methods with the correction. We evaluate this correction on a single 3-timePlex run in Fig. 4a using the mixed-species standards to benchmark the change in quantitative accuracy, and indeed observe improved accuracy. Next, we sought to evaluate the quantitative performance of both timePlex implementations (‘separate emitters’ and ‘single emitter, merged flow’). The data indicate accurate precursor-level quantitation for both approaches, as shown in Fig. 4b.

**Figure 4.**
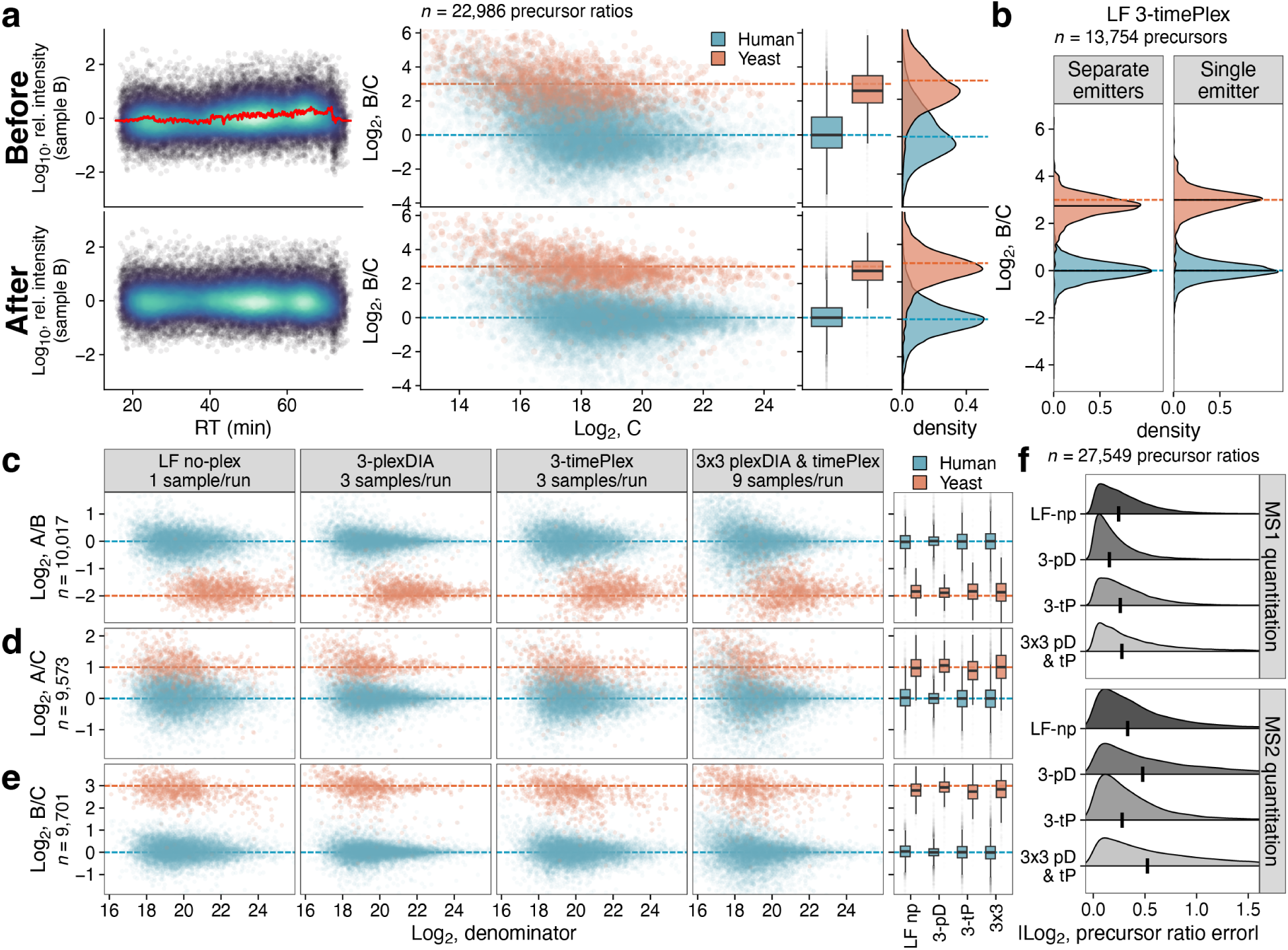
Benchmarking timePlex quantitative accuracy. **a** Precursor intensities may vary across the gradient in timePlex. This change is systematic and correctable as shown in quantitative assessments before and after the correction was applied. The relative intensity panels correspond to one timePlex channel in a single run and the quantitative ratio panels compare that timePlex channel to another channel in that same run, in this case Sample B and C are shown for yeast (orange) and human (blue) precursors. All quantitation is from MS1 precursor intensities measured by Exploris 240. **b** Density plots for MS1 ratios of human and yeast precursors in common between samples B and C (*n*=13,754) are shown for both timePlex implementations to assess quantitative accuracy. Black horizontal lines mark the median of each respective distribution. **c-e** MS1 quantitation is shown for precursor ratios in common across all four methods and shown as scatter and box plots. Scatter plots show the 0.5th–99.5th percentiles of values; box plots reflect all ratios. **f** Comparisons of MS1 and MS2-level quantitation are shown for intersected precursor ratios across methods. The following abbreviations are used: LF-np: LF no-plex, 3-pD: 3-plexDIA, 3-tP: 3-timePlex, and 3×3 pD & tP: 3×3 plexDIA & timePlex. The absolute-values of the errors between expected and observed fold-changes for each precursor ratio are shown as distributions with black vertical lines indicating the median.

### Benchmarking timePlex quantitative accuracy

We continued to use these mixed species standards to benchmark the quantitative performance of 3-plexDIA, 3-timePlex, and combinatorial 3×3 timePlex & plexDIA to LF no-plex data using the ‘separate emitters’ implementation. The number of quantified precursor ratios between sample pairs for the four methods is shown in Extended Data Fig. 7a-c; despite the large throughput differences, a similar number of precursor ratios were quantified between methods with the exception of the combinatorial 3×3 timePlex & plexDIA (9-plex), which resulted in an approximately 30% reduction. To fairly compare quantitative accuracy between methods, only the subset of precursor ratios quantified across all methods was used for accuracy benchmarking, Fig. 4c-e. All multiplexed approaches produced comparable median quantitative accuracy to LF no-plex data, with a reduction in precision observed for the 9-plex combination of 3×3 timePlex & plexDIA, which further computational modeling of retention time and intensity based effects may improve (Extended Data Fig. 6).

We sought to evaluate whether quantitative accuracy of 3×3 plexDIA & timePlex varied at high density regions of complexity. To do this a combinatorially multiplexed run of 3×3 plexDIA & timePlex was benchmarked for each time channel with respect to retention time; we observed no complexity-dependent changes in quantitative accuracy. This suggests combinatorial plexDIA & timePlex quantitation remains accurate and relatively invariant to differences in temporal complexity, Extended Data Fig. 8. We summarize quantitative accuracy investigations at MS1 and MS2-levels by computing absolute errors between observed and expected fold-changes for intersected precursor ratios (*n*=27,549) quantified across all methods. Quantitative accuracy remained similar across all methods at MS1 and MS2-levels, with MS1 quantitation being the most accurate in each case, Fig. 4f.

### Realizing 27-plex multiplexing with 9×3 plexDIA & timePlex

Recent developments in mass-tag engineering have produced tags that improve spectral quality and enable 9-plexDIA^32^. We paired these novel mass tags (termed ‘PSMtag’) with 3-timePlex to enable a combinatorial 27-plex. Using 25 minutes of active chromatography per time channel, we exceeded 500 samples/day. The total duration per run was 76 minutes, approximately 23.5 minutes of which was sample-loading and 52.5 minutes was data acquisition, column washing, and equilibration.

We benchmarked this 27-plex with 1 ng mixed-species proteomic digests on a Thermo Orbitrap Exploris 480 with FAIMS, the compositions for which are shown in Fig. 5a. As a first evaluation of the raw data, we display the extracted ion chromatograms for select human and yeast precursors in Fig. 5b. We observed differential abundance of yeast precursors at their expected spike-in amounts, while human precursors remained similarly abundant across all mass and time channels. In this data acquisition, we again controlled for the timePlex setup and acquired triplicates of all four samples with 9-plexDIA and combinatorial 9×3 plexDIA & timePlex. The number of precursors and proteins identified is shown in Fig. 5c, averaging approximately 12,400 precursors/sample and 3,300 proteins/sample for 9-plexDIA, and 9,200 precursors/sample and 2,800 proteins/sample for combinatorial 9×3 plexDIA & timePlex. The mixed species spike-ins were used to benchmark precursor-level quantitative accuracy for all methods. The median quantification of precursor ratios was found to be accurate for the 9-plex and combinatorial 27-plex, with a modest decrease in quantitative accuracy and precision with combinatorial multiplexing Fig. 5d. These results demonstrate the ability to generate quantitative data with 27-plex DIA using combinatorial plexDIA and timePlex.

**Figure 5.**
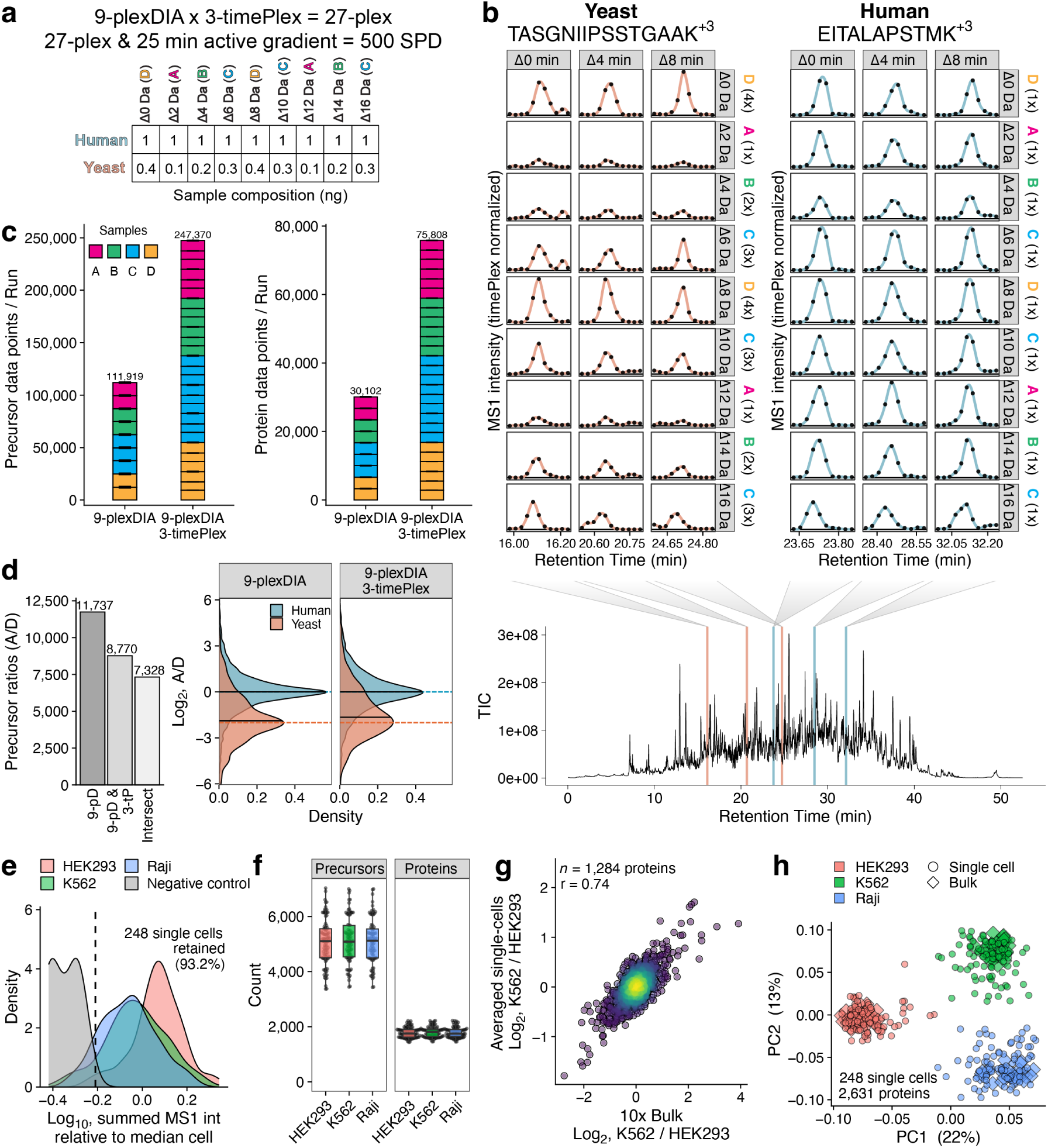
Combining 9-plexDIA with 3-timePlex enables 27-plex. **a** Proteomic compositions of samples A-D consisted of 1 ng human digest and variable amounts of mixed yeast digest (0.1-0.4 ng) and acquired on a Thermo Orbitrap Exploris 480 mass spectrometer. These samples were labeled with PSMtags and pooled to enable 9-plexDIA. Samples were acquired at a rate exceeding 500 samples/day with combinatorial 9×3 plexDIA & timePlex. **b** Extracted ion chromatograms for yeast and human precursors from the 9×3 plexDIA & timePlex data. **c** Number of sample-specific precursors and proteins identified per run for 9-plexDIA and 9×3 plexDIA & timePlex. **d** Bar plots show the number of precursor ratios quantified across 9-plexDIA and 9×3 plexDIA & timePlex. The intersecting precursors across methods were used to benchmark quantitative accuracy at MS2-level, shown as density plots of the measured ratios. **e** Density plots for summed MS1 intensity for single cells and negative controls relative to the median single-cell intensity. The data were acquired at *>*500 SPD using Orbitrap Astral Zoom with 9×3 plexDIA & timePlex. **F** Number of precursors and proteins identified per single cell. **g** Scatter plot of protein fold changes between averaged single-cells and 10-cell bulk standards for the subset of proteins quantified in *≥*50% of single cells. **h** PCA of single cells and 10-cell bulk standards of HEK293, K562, and Raji cells.

Given that the combination of timePlex & plexDIA enables similar quantitative accuracy at combinatorially higher throughput, we sought to evaluate its potential in maintaining high quality quantitation while scaling up the throughput of single-cell proteomics. Single cells from K562, Raji, and HEK293 human cell-lines were prepared by nPOP^56,57^ using PSMtag to isotopically label single cells to be pooled into single-cell plexDIA sets. The 9-plexDIA sets were acquired with 3-timePlex using approximately 3 minute offsets on a Thermo Orbitrap Astral Zoom mass spectrometer. Single cells were separable from negative controls which underwent the same preparation but lacked cellular material, Fig. 5e. Searching these data with JMod quantified a median of 5,102 precursors and 1,758 proteins per cell without using match between runs, Fig. 5f.

To benchmark the reliability of our single-cell measurements, bulk lysate was diluted to 10-cell amounts per channel and acquired as combinatorial 9-plexDIA & 3-timePlex. Fold-changes between cell-types indicate quantitative agreement between the relative protein abundances averaged across single cells and the diluted bulk lysate, with a Pearson correlation of 0.74 for the subset of proteins that have *≥*50% data completeness, Fig. 5g. Dimensionality reduction by PCA of 10x diluted bulk and single-cell data from 9×3 plexDIA & timePlex is shown in Fig. 5h. PC1 and PC2 produced clear separation between cell-types and co-clustering of single-cell 9×3 plexDIA & timePlex data and 10x bulk cell lysates. These results highlight the potential for combinatorial multiplexing of timePlex and plexDIA to increase throughput of single-cell proteomics.

### Dual-timePlex analyzes *>*1,200 samples in 24 hrs

While combinatorial multiplexing with timePlex and plexDIA achieves a 27-plex analysis with DIA, throughput was limited by overheads associated with sample loading, column washing, and equilibration. We sought to reduce these overheads by implementing a ‘dual-timePlex’ approach analogous to dual-column or dual-trap methods previously developed^2,3^. This allows for simultaneous sample-loading onto one timePlex set while the other timePlex set actively separates samples, thus reducing overheads. A schematic for the dual-timePlex approach can be found in Extended Data Fig. 9. In short, two LC-systems were utilized where one provided isocratic flow for equilibration and sample-loading and the other delivered the active gradient and wash. We visualize the conceptual operation of dual-timePlex in Fig. 6a, whereby one timePlex set (Set A) is actively separating 9×3 plexDIA & timePlex samples while the other set (Set B) is loading, alternating between the two sets to reduce overheads. A custom 3D-printed setup was engineered to adjust the emitter tips and hold the dual-timePlex configuration in place, as shown in Fig. 6b.

**Figure 6.**
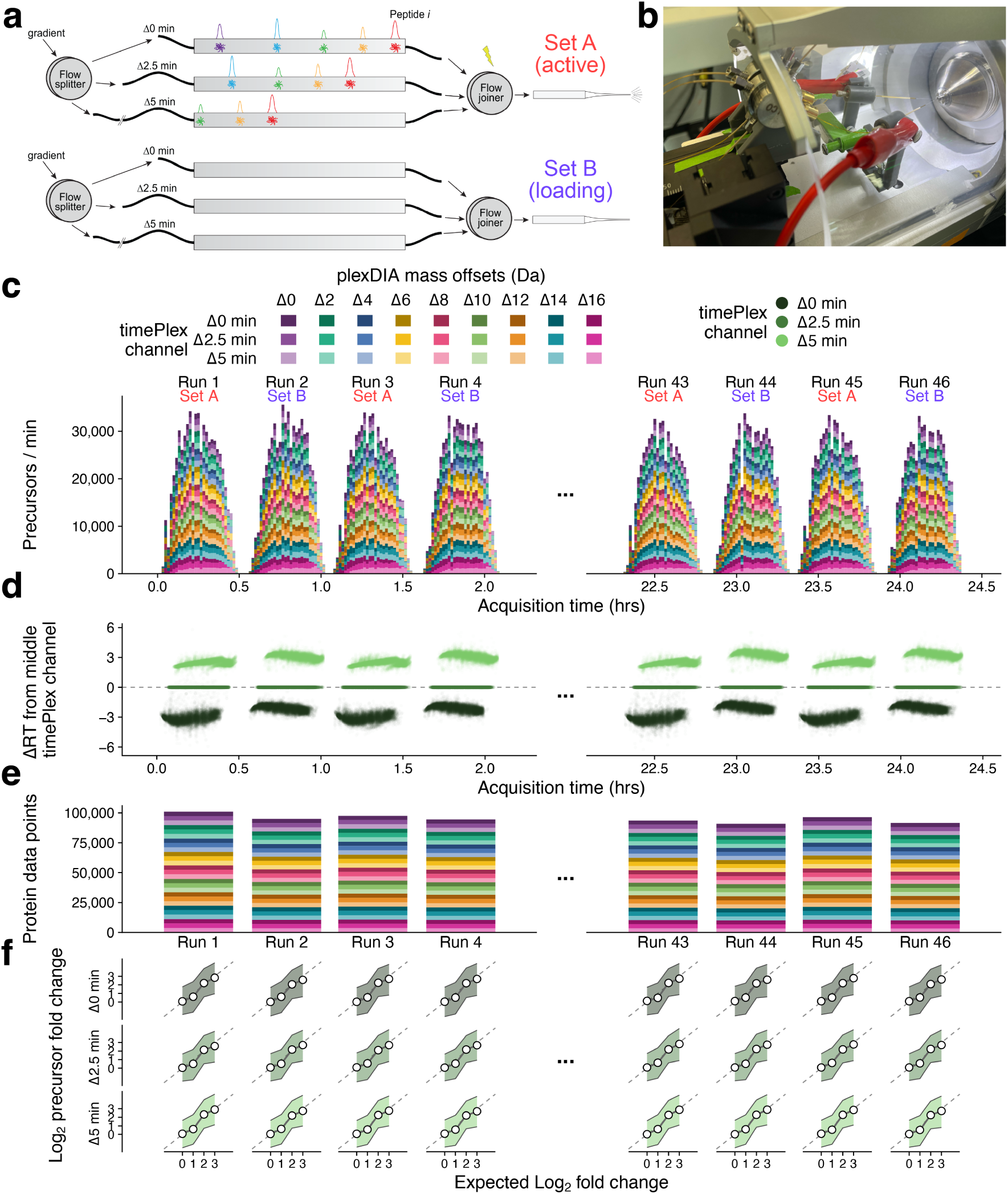
Exceeding 1,200 SPD with 9×3 plexDIA & dual-timePlex. **a** Conceptual visualization of dual-timePlex operation. One timePlex set (e.g. Set A) actively separates samples while another set (e.g. Set B) simultaneously loads samples in preparation for eventual switching. **b** Image of the dual-timePlex setup implemented in this work. **c** Histograms of the number of precursors identified per minute across 24 hours of operation at *>*1,200 SPD, showing the first four and last four runs. **d** Evaluation of consistency of RT-offsets between timePlex channels over 24 hours of operation. Each point represents a precursor RT from a given timePlex channel and its difference from the middle timePlex channel is shown on the y-axis. **e** Number of proteins identified per sample in each 9×3 plexDIA & timePlex run, shown as the sum for each run. **f** Precursor quantitation for each timePlex channel is shown for each run where the y-axis corresponds to the observed fold-change and x-axis is the theoretical expected fold-change based on the composition yeast spike-in. Each dot represents the median precursor fold-change, the shaded ribbon corresponds to the interquartile range for each spike-in amount, and the dashed line represents a slope of 1.

This approach combined with 9×3 plexDIA & timePlex and microflow separations (5 *µ*L/min) achieved throughput of *>*1,200 samples per day (SPD) – quantifying over 4 million protein data points in a day – using approximately 23 minutes of active chromatography per sample. To evaluate the depth of coverage, robustness, and quantitative accuracy of this workflow, we analyzed 100 ng samples composed of human and yeast spike-ins using an Orbitrap Astral mass spectrometer for 24 hours, Fig. 6. The number of precursors identified per run is shown as a stacked histogram in Fig. 6c for the first four and last four runs of the 24 hour acquisition. In addition, the column performance was consistent across all columns with a median FWHM of 2.5s across precursors in common Extended Data Fig. 10a; likewise, the number of precursor and protein identifications between different columns of Set A and Set B was found to be comparable, with an average of approximately 21,000 precursors and 3,500 proteins per sample for Set A and Set B, as shown in Extended Data Fig. 10b-c, which held for 24 hours (Extended Data Fig. 10d-e). These experiments achieved approximately 2-fold higher depth of coverage at 50-fold higher throughput than previous benchmarks at 5 *µ*L/min^58^. The higher depth of coverage likely owed to the more sensitive Orbitrap Astral MS instrument. Furthermore, the time-offsets of these precursors across time channels were found to be separable and consistent with each respective set for the 24 hour period, as shown in Fig. 6d; the set-to-set differences correspond to inherent differences between the encoded time-offsets of the two timePlex sets. The number of proteins identified per run is shown in Fig. 6e, on average identifying approximately 95,000 protein data points/run.

Generating quantitatively accurate data at a throughput exceeding hundreds of samples per day is a known challenge with DIA proteomics. Therefore, we sought to evaluate the quantitative accuracy of combinatorial multiplexed data at *>*1,200 SPD, and its consistency over 24 hours of operation. We found that the median quantitation remained accurate within each timePlex channel at up to 8-fold spike-in ratios and throughout the 24 hours of acquisition, as shown in Fig. 6f. Together, these data suggest combinatorial plexDIA & timePlex multiplexing can robustly operate at *>*1,200 SPD for 24 hours, enabling accurate quantification of approximately 21,000 precursors and 3,500 proteins per sample.

## Discussion

Here we demonstrate that multiplexing in two domains (mass and time) enables combinatorially increased sample-throughput. Time-based multiplexing (‘timePlex’) was developed in this work to enable this accelerated throughput when paired with plexDIA, and was specifically demonstrated with 3×3 plexDIA & timePlex to enable a 9-plex, and 9×3 plexDIA & timePlex to enable a 27-plex. Because multiplexing in the time-domain is label-free, the technique applies to analytes that cannot be easily labeled, such as metabolites.

Sample-throughput can also be increased without multiplexing, for example by compressing the LC-separations of each sample. Indeed, recent work has demonstrated the identification of *>*7,000 protein groups using the Evosep Eno LC system on an Orbitrap-Astral Zoom mass spectrometer at 500 SPD without multiplexing^59^. Benchmarking this 500 SPD workflow indicated significant challenges to achieving quantitative accuracy because of too few points per peak even with the fastest duty cycles^59^. In contrast, the protein ratios measured with our 1,200 SPD methods center around the corresponding mixing ratios (Fig. 6) because timePlex enables this throughput with longer separation times accommodating more points across elution peaks. This makes timePlex approaches a promising direction for maintaining sufficient peak sampling and quantitative accuracy at very higher throughput rates that will complement existing short gradient workflows affording easier implementations at lower throughput rates.

In this work, timePlex was demonstrated with two accessible implementations, which utilized separate RP-HPLC columns. Future implementations may be improved, for example by implementing isocratic separation and continuous data acquisition. Such implementation can allow using a single column and limiting LC-associated overheads. Another option is using capillary electrophoresis (CE), analogous to metabolomics implementations using multi-segment injections for simultaneous acquisition of multiple samples per run through staggered and overlapped separations^41,60^; as such, high-timePlex proteomics may be readily implemented through multi-segment CE approaches. Another approach could be size-exclusion chromatography (SEC) with isocratic flow as this could potentially facilitate high-timePlex from a single column. While such approaches remain to be explored, a related but more limited implementation of 2-timePlex was reported by another laboratory after the release of our initial preprint^61^. That implementation lacked benchmarking of quantitative accuracy against known ratios and required a dedicated UHPLC per column, but it nonetheless supports the feasibility of broader adoption of timePlex.

While 27-plex analysis by DIA was achieved through combinatorial multiplexing of plexDIA and timePlex, the timePlex implementation in this work involved some drawbacks that future work can mitigate. In particular, 1) intensity variation over the course of the gradient needed correction to reduce adverse effects on quantitation (Fig. 4a), and 2) current implementations resulted in higher overall flow rates, which reduced sensitivity and depth of coverage. Regarding the former, differences in desolvation and ion suppression over the course of the gradient may result in dynamically changing ion intensities over time, among other factors which we have not fully characterized. Correcting these rapid changes becomes more difficult for time regions with sparse data. However, we found this effect remained largely correctable as observed in our benchmarking results. Regarding the latter, timePlex coverage will be improved through the use of lower flow rates and narrower inner diameter columns to offset the time-overheads associated with lower flow rates. While the timePlex implementations in their current form may reduce ion intensities and thus proteomic coverage at 200 nL/min/column, lower flow rates were found to make the intensities equivalent (Extended Data Fig. 2). Furthermore, the timePlex data type itself, as benchmarked in this work by controlling for the timePlex setup, indicate comparable quality to non-multiplexed DIA with quantitative accuracy that was found to be invariant to changes in spectral complexity that occur throughout the run (Extended Data Fig. 8).

The throughput demonstrated by 9×3 plexDIA & dual-timePlex (*>*1,200 SPD) has only been achieved so far using isobaric mass tags^57,62^, which have well characterized limitations^63,64^. We expect the coverage will substantially improve when implemented with lower flow rates, longer columns with smaller inner diameters and unions that introduce lower post-column dead volumes. The complexity of highly multiplexed DIA data can be better modeled by searching data with spectral libraries whose fragmentation patterns more accurately reflect empirically observed spectra, as achieved through transfer-learning approaches^52^. In addition, we expect implementations in the near-future utilizing shorter chromatographic separations and higher plex will enable throughputs exceeding 5,000 SPD. Realizing this promise of higher-multiplexed DIA necessitates the coordinated development of data interpretation approaches for highly parallelized mass spectra such as JMod^44^, and continued advances in LC-separation equipment, methods, and mass spectrometry instrumentation. The convergence of these advances will reduce the time and costs associated with LC-MS proteomics and support the generation of larger-scale proteomic datasets at high depth and accuracy.

## Methods

### timePlex implementation

timePlex was implemented in this work using a Dionex UltiMate 3000 UHPLC. Schematics and details for sample loading are shown in Supplementary Fig. 2. Flow from the NC pump was split using a ZenFit tee (700011518, Waters), and nanoViper™ transfer lines ranging from 20-50 *µ*m ID and 150 mm - 950 mm were used in combination to encode time offsets between samples.

Sample-loading occurred in the following way: The first sample was injected with valves in the 1 2 and 1 2 position for the first and second valves. At this stage, flow is pushed through the sample loop into port 1 of Valve 2, which flows out of Port 2 and into Port 8 of Valve 1, and out of Port 7 to ‘Column 1’. The flows which were split from the NC pump push against plugs; as a result, all net flow is directed through the sample loop. At this step, only ‘Column 1’ is actively receiving flow. To prevent over-pressurization, the flow rate at the step is set to 300 nL/min. Immediately after injection, the sample is run isocratically for 7 minutes with 2% ACN and 0.1% formic acid, for the sole-purpose of pushing the sample out of the loop and onto the first column. This process takes approximately 10 minutes total.

The second column is loaded in a similar way, except Valve 2 is switched to the 1 10 position, directing the second sample onto ‘Column 2’. This process also takes approximately 10 minutes total. In the last 10 seconds, the flow rate is increased to 600 nL/min in anticipation of switching the valves which occurs during the loading of the third column where all three columns are receiving flow.

The third column is loaded by switching the positions of both valves to 1 10 and 1 2 positions, respectively for the first and second valves. At this point, all three columns receive flow at a total rate from the pump of 600 nL/min. Therefore, the flow rate across each column is approximately 200 nL/min although differences in the resistances of the columns may lead to unequal flow rate across each column, and thus the encoded time-differential. As a result, the volume differences which encode the time-offsets may need to be adjusted to achieve the desired time-differential between timePlexes. The gradient can start immediately when loading the third sample. Additionally, because the third sample will receive its gradient through the sample loop, which in this case holds a 1 *µ*L volume, it was most-efficient to have this column be the latest eluting one. In this work, transfer lines of specific lengths and inner diameters were used to encode volumetric and thus time differences between timePlex channels. The volume of each transfer line was estimated by calculating the volume of the cylinder based on each transfer line’s inner diameter and length. To encode 4 minute offsets (800 nL offsets at 200 nL/min/column), a 50 *µ*m x 350 mm transfer line (Thermo Scientific, nanoViper 6041.5540) was connected to a 20 *µ*m x 350 mm transfer line (Thermo Scientific, nanoViper 6041.5240) as a general starting point, removing or adding transfer lines of known volume as necessary to tune the timePlex channel offsets; such tuning may be necessary if resistances (and thus flow) are unequal across columns. All samples, including non-timePlex data acquisitions (such as LF no-plex and plexDIA), were acquired with the timePlex setup so we could directly evaluate the impact on timePlex data on downstream interpretation rather than the combination of interpretation and implementation. This enabled direct evaluation of timePlex data quality.

### Differences between single and multi-emitter approaches

Both timePlex implementations demonstrated in this work utilize the same sample-loading scheme and a Nanospray Flex™ Ion Source on an Orbitrap Exploris 240. The difference between the multiemitter (separate flow) approach and the single-emitter (joined flow) approach only pertains to how the flow is treated post-column separation and how the ESI occurs.

In the case of the multiple emitters approach, voltage (+2.2 kV) was applied from the MS source to two columns via an alligator clip, and an auxiliary power supply (PS350, Stanford Research Systems) applied voltage (+2.2 kV) to the third column. A PEEK cone, kindly provided by IonOpticks, held the columns in place, pointing spray at the mass-spec inlet of an Exploris 240.

For the merged flow approach, flow out of each *µ*PAC column was merged through a 50 *µ*m bore cross (C360TS62, VICI Valco) to an emitter tip (TIP360002010-10-5, MS WIL). Utilizing narrow-bore crosses for merging flow post-column mitigated peak broadening. Voltage (+2.2 kV) was applied to the 50 *µ*m bore cross via alligator clip.

### Dual-timePlex implementation

Dual-timePlex was enabled through the use of two LC systems, specifically a Dionex UltiMate 3000 UHPLC and a Thermo Scientific Vanquish Neo UHPLC. Samples were loaded using isocratic flow on the Dionex UltiMate 3000 at 24% Mobile Phase B (100% ACN & 0.1% formic acid) to account for the increased hydrophobicity of samples labeled with PSMtag. The column compartment 10-port valves were utilized to enable switching between timePlex sets, as shown in Extended Data Fig. 9. Samples were delivered to their appropriate column through a multi-position 8-port high pressure selector valve (VICI Valco, C85X-6678EUTA), with timed switching encoded for each subsequent injection.

The approach utilized a merged-flow approach in which each timePlex set consisted of three custom 15 cm x 150 µm ID columns with 1/16” fittings (IonOpticks) which merged into a 150 micron bore cross union (VICI Valco, ZX1XCS6). The cross was connected to a 20 µm x 6 cm transfer line ending in a silica to ferrule connection to a 20 µm x 50 mm Constant i.d. Etched-Tip (CET) ESI Emitter (MicrOmics Technologies, EMCA). A custom 3D-printed mount held the union and emitter in place, allowing the emitter’s position and angle to be adjusted while remaining stable during acquisition. Voltage (+2.6 kV) was applied to the union that connected the transfer line to the emitter. Voltage-switching was coordinated automatically through the method editor relay on the Dionex UltiMate 3000. This relay triggered switching on the high-voltage relay, thus directing voltage to the appropriate timePlex set.

The LC-gradient was delivered by the Thermo Scientific Vanquish Neo UHPLC in which its activation was triggered by a relay on the Dionex UltiMate 3000 UHPLC. The coordination of sample-loading and gradient delivery was facilitated by each LC-system in communication with its own respective Thermo Scientific Xcalibur queue, the Dionex consisted only of sequential sample loading while also serving as the trigger for initiation of the gradient on the Vanquish Neo and MS-acquisition. A diagram of the setup can be found in Extended Data Fig. 9 and instrument methods for sample loading and gradient delivery and MS acquisition can be found in our data repository.

### MS data acquisition

All samples were resuspended in 0.015% n-Dodecyl *β*-d-maltoside (DDM)^65^ with 0.1% formic acid, with the exception of PSMtag-labeled samples which were resuspended in 0.03% DDM with 24% acetonitrile and 0.1% formic acid to account for the added hydrophobicity introduced by the tag. Each benchmarking sample, which consisted of 8 ng human digest and variable yeast digest, was acquired with the timePlex multiplexing setups shown in Fig. 2a, whether it was timePlexed or non-timePlexed. This means LF no-plex and 3-plexDIA benchmarks were acquired with the same setup as the timePlex experiments (3 columns active at a time).

Benchmarking data for 3×3 plexDIA and timePlex were acquired on a Thermo Orbitrap Exploris 240 orbitrap mass spectrometer using either 25 cm x 75 *µ*m IonOpticks columns (AUR3-25075C18) or 50 cm *µ*PAC™ Neo columns (COL-NANO050NEOB) with a Dionex UltiMate 3000. 8 ng LF samples were acquired in DIA mode with an MS1 scan at 120k resolving power and 300% AGC conducted every 1.3 seconds. The MS2 duty cycle consisted of 24 variable length windows covering 378 - 1356 *m/z* for LF and 478-1356 *m/z* for mTRAQ-labeled peptides. The MS2 windows were designed to equally distribute the ion intensities over the MS1 *m/z* range.

Benchmarking data for 9×3 plexDIA and timePlex were acquired on a Thermo Orbitrap Exploris 480 mass spectrometer with FAIMS Pro (CV=-50 and 5.2 L/min carrier gas) with 25 cm x 75 *µ*m IonOpticks columns (AUR3-25075C18) and Dionex UltiMate 3000 HPLC.

Single-cell data for 9×3 plexDIA and timePlex were acquired on a Thermo Orbitrap Astral Zoom mass spectrometer with FAIMS Pro (CV=-50 and 4 L/min carrier gas) with 25 cm x 75 *µ*m IonOpticks columns (AUR3-25075C18) and Dionex UltiMate 3000 HPLC.

Bulk benchmarking data for 9×3 plexDIA and dual-timePlex were acquired on a Thermo Orbitrap Astral mass spectrometer with FAIMS Pro (CV=-50 and 4 L/min carrier gas) with custom 15 cm x 150 µm ID columns with 1/16” fittings (IonOpticks) using a Dionex UltiMate 3000 HPLC for sample loading and a Thermo Vanquish Neo UHPLC for gradient delivery.

### LF benchmarking

For LF samples used in the 3×3 plexDIA and timePlex benchmarks, the MS2 windows were as follows: 378-387, 386.5-393, 392.5-401.5, 401-409.5, 409-418, 417.5-427.5, 427-438.5, 438-448.5, 448-460.5, 460-473.5, 473-487, 486.5-501.5, 501-517, 516.5-536.5, 536-558.5, 558-580.5, 580-607, 606.5-639, 638.5-675.5, 675-723.5, 723-784.5, 784-867, 866.5-986, and 985.5-1356 *m/z*.

Gradient for IonOpticks columns ‘separate emitters’ approach: 600 nL/min ramping from 2% B to 5.8% 2.25 minute into the gradient, 8.8% 6.3 minutes into gradient, 11.5% (11.25 min), 18.5% (31 min), 25.5% (45 min), 34.2% (54.9 min), 37.7% (56.25 min), 45% (60 min), 48% (68 min), 95% (68.2 min), 95% (72 min), 2% (72.1 min), maintaining 2% B until end (90 min). The full LC-MS method was 90 minutes.

Gradient for *µ*PAC columns ‘single emitter’ approach: 600 nL/min ramping from 2.5% B to 4% 1 minute into the gradient, 8% 5.5 minutes into gradient, 15% (26 min), 18.5% (31.2 min), 25% (37.4 min), 32% (41 min), 34% (51.5 min), 95% (52 min), 95% (58 min), 2% (64 min), maintaining 2% B until end (75 min), with the last 10 minutes run at 1,200 nL/min. The full LC-MS method was 75 minutes.

### mTRAQ-labeled benchmarks

For mTRAQ-labeled benchmarks, the MS2 windows were as follows: 478-485.5, 485-493, 492.5-501.5, 501-510.5, 510-518, 517.5-526.5, 526-535.5, 535-544.5, 544-553, 552.5-563, 562.5-574.5, 574-586.5, 586-598.5, 598-612.5, 612-627, 626.5-643.5, 643-662, 661.5-683, 682.5-707.5, 707-739, 738.5-778, 777.5-834, 833.5-940, and 939.5-1356 *m/z*.

Gradient for IonOpticks columns ‘separate emitters’ approach: 600 nL/min ramping from 2% B to 6.5% 2.25 minute into the gradient, 9.5% 6.3 minutes into gradient, 12% (11.25 min), 19.4% (31 min), 26.4% (45 min), 35.1% (54.9 min), 46% (60 min), 49% (68 min), 95% (68.2 min), 95% (72 min), 2% (72.1 min), maintaining 2% B until end (90 min). The full LC-MS method was 90 minutes.

Gradient for *µ*PAC columns ‘single emitter’ approach: 600 nL/min ramping from 4% B to 8% 5.5 minute into the gradient, 15% (26 min), 19.5% (31.2 min), 26% (37.4 min), 33.3% (41 min), 36.5% (51.5 min), 95% (52 min), 95% (58 min), 2% (58.1 min), maintaining 2% B until end (75 min), with the last 10 minutes run at 1,200 nL/min.

### PSMtag-labeled benchmarks

For PSMtag-labeled benchmarks, FAIMS was utilized with CV = -50 with an MS1 scan at 180k resolving power (500% AGC) performed every 1.3 seconds. The MS2 scans were performed at 45k (1000% AGC) resolving power with the following windows: 500-538.5, 538-566.5, 566-590.5, 590-611.5, 611-633.5, 633-649.5, 649-663.5, 663-674.5, 674-685.5, 685-700.5, 700-716.5, 716-728.5, 728-743.5, 743-754.5, 754-771.5, 771-789.5, 789-816.5, 816-847.5, 847-893.5, 893-1000 *m/z*.

The gradient was as follows: 600 nL/min ramping from 24% B to 30.4% 0.15 minutes into the gradient, 49.6% 20.4 minutes into gradient, 60% (25 min), 62% (31.5 min), 95% (32 min), 95% (37 min), 24% (37.1 min), and at 47.1 min the flow rate is increased to 1 *µ*L/min until 50 minutes, and then lowered to 300 nL/min until the end (52.5 minutes).

### Dual-timePlex PSMtag-labeled benchmarks

For PSMtag-labeled benchmarks, an MS1 scan at 240k resolving power (600% AGC) was performed every 0.6 seconds, scan range = 498-1002 *m/z*, and max IT = 20 ms. MS2 scans were performed using 2-Th windows spanning 500-1000 *m/z* range using the Astral detector (500% AGC), NCE = 24, max IT = 3 ms, and scan range = 180-1800 *m/z*

To load samples for Set A while delivering gradient to Set B, the 10-port 2-position valves of the Dionex UltiMate 3000 were in 1 2 & 1 2 valve positions, using the selector valve to direct each sample to its appropriate column. Alternatively, to load samples for Set B while delivering gradient to Set A, the Dionex UltiMate 3000 was in 1 10 & 1 10 valve positions. Each sample was sequentially loaded for 8.7 minutes at a flow rate of 1,200 nL/min.

The gradient was as follows: 5 *µ*L/min throughout, ramping from 30% B (Mobile Phase B: 80% ACN & 0.1% formic acid) at the start of the gradient to 50% 17.6 minutes into the gradient, 60% (21.6 min), 62% (26.6 min), 99% (26.65 min), 99% (28.65 min), 1% (28.7 min), and held at 1% until the end of the run (29.25 minutes). The gradient was delivered to the appropriate timePlex set by using a 6-port 2-position valve in the column compartment of the Thermo Vanquish Neo UHPLC.

### plexDIA and timePlex single-cells

For plexDIA and timePlex single-cell acquisition on the Thermo Orbitrap Astral Zoom, MS1 data was acquired every 0.6 seconds using enhanced dynamic range enabled^66^ which divided the 498 - 1002 m/z range into 6 accumulations to improve MS1 sensitivity. Additional MS1 settings were as follows: AGC max = 500%, max IT = 80 ms, resolving power = 240k, and profile mode enabled. MS2 scans split the range into 40 windows with an average size of 13 Th and 0.5 Th overlapping regions. The following additional settings were used: NCE = 24, scan range = 180-1800, pre-accumulation enabled, Max IT = 20 ms, AGC = ‘Standard’. The gradient proceeded as described in the ‘PSMtag-labeled benchmarks’ section.

### Spectral library generation with DIA-NN

Data was generated to create empirical libraries for searching the human and yeast benchmarking data. The library-generating data consisted of 250 ng injections of label-free and mTRAQ-labeled human (K562) and yeast standards, acquired on the Exploris 240 using 25 cm x 75 *µ*m IonOpticks columns (AUR3-25075C18) and a Dionex UltiMate 3000 HPLC. The data was searched with DIANN version 1.9^15^ using a predicted spectral library. To generate an empirical library for searching single-cells, 30-cell injections of PSMtag-labeled Raji, K562, and HEK293 protein digests were similarly searched starting with a predicted library created as previously described^67^.

### Retention time prediction and transfer learning

Empirical libraries output from DIA-NN^15^, which were generated from label-free and mTRAQ-labeled data, were used to train a model to predict peptide iRTs. Precursors from the empirical libraries were collapsed to peptide-level, averaging the iRT between charge-states, if there were any reported differences. The peptides were one-hot encoded as a 30x20 array for each peptide, where 30 is the maximum length for a peptide and 20 is the number of possible amino acids. A hybrid convolutional and fully connected feedforward neural network was trained to predict peptide RTs using TensorFlow. The model consisted of four 1D convolutional layers, with the number of filters doubling at each successive layer: 32, 64, 128, and 256. Each layer used a kernel size of 4. No pooling was applied after the first two convolutional layers. Max pooling with a pool size of 2 was applied after each of the final two convolutional layers. The output of the final convolutional layer was flattened and passed into a fully connected feedforward neural network with five layers consisting of 1000, 256, 128, 64, and 32 nodes, respectively. All layers used the ‘Swish’ activation function^68^, except for the final output layer, which used a linear activation to produce the final RT value. An ensemble of five models was created and used for transfer learning; predictions were the average of the five model outputs. The models were trained with early stopping (monitor=’val loss’, patience = 20, restore best weights=True), epochs = 150, batch size = 32, validation split = 0.1, with Adam^69^ as the optimizer (learning rate=0.001) and mean absolute error as the loss function.

Transfer learning was applied to fine-tune library RTs. This involved re-initializing the five models and training on new data stemming from the initial search which occurs in JMod for *m/z* and RT alignment. The transfer learning occurs under the following parameters: early stopping was enabled (monitor=’val loss’, patience = 10, restore best weights=True), epochs = 50, batch size=24, optimizer=’Adam’ (learning rate=0.0001), loss = ‘mae’, validation split = 0.1. The average RT output across the five models is used as the final RT prediction in the library. Finally, the RT predictions from transfer learning are compared to the original library RTs using the validation set of data. The strategy with the greatest RT accuracy is used for the full-search in JMod.

### Searching with JMod

JMod is an open source proteomics search software, which leverages joint-modeling of spectra to enable interpretation of proteomics data acquired by DIA^44^. In this work we developed a version of JMod to enable searching of timePlex and combinatorially multiplexed plexDIA and timePlex data (JMod v2.0.0 timePlex beta). First, a preliminary search is used for calibrating precursor *m/z* errors, retention time fine-tuning, and optimizing the *m/z* and retention time tolerances. This is performed using peppy sage an internally developed Python library that was forked from Sage^70^, reporting up to 15 candidate precursors per scan per multiplexing channel. Spectral angle and Scribe scores are computed for these identifications. This first search does not utilize any retention time information and, due to JMod’s approach, can identify multiple instances of the same precursor. For *n*-timePlex, identified peptides found in exactly *n* clusters are extracted, using the Scribe score cutoff that maximizes the number of such peptides. These are assumed to have one identification for each column, which are then assigned to each timePlex channel based on their temporal elution order. The time differences encoded by the timePlex setup remain relatively constant or vary smoothly across the gradient. Therefore, correctly matched precursor sets will have systematic offsets or potentially a dynamically changing offset that JMod models, which may occur due to dynamically changing column resistances. Importantly, JMod does not require the same offset to be encoded between different pairs of successive timePlex-channels and does not require the user to specify the offset; it is modeled automatically. However, the consistency between each pair of columns allows us to further filter our set of precursors to those that agree on the time encoding. This provides a set of high-confidence identifications across the gradient which are separated into different timePlexes. A LOWESS regression is fit to each individual timePlex channel to align the library retention times to the observed data. In addition, the peptides from each timePlex channel are used to independently fine-tune a retention time prediction model. LOWESS regressions are also fit to the model’s predicted retention times vs the observed retention times for each timePlex to convert them to observed retention times.

For each timePlex channel, the differences between these fits and the observed retention times of held-out peptides (10%, withheld from fitting and fine-tuning) are used to calculate retention time errors. Cumulative distributions of the absolute errors are calculated for both the original library retention time errors and the newly predicted retention time errors. The cumulative distribution with the largest area under the curve is selected as the best fit Supplementary Fig. 4, allowing each channel to use its best-fit retention times, which may be either the original library or the fine-tuned predictions. Then, to calculate the optimal retention time tolerance, a two-component, zero-mean Gaussian mixture model is fit to the held-out retention time errors for each channel. Each timePlex channel’s retention time tolerance is set to *±*4 standard deviations of the narrower component, as this narrower component likely reflects correct identifications.

Once a best model is selected, JMod duplicates the library for each timePlex channel. Each copy will have an updated retention time based on the model selected for that channel (its LOWESS-fit original retention times or its fine-tuned predicted retention times). These tolerances are also made to be non-overlapping between adjacent timePlex channels; if adjacent tolerance windows were to overlap, both would be reduced to at most half of the time difference between the timePlex channels. This ensures that the same precursor in each timePlex channel will be considered independent of the other channels, leading to column and sample-specific identifications. Once the library is created, and each channel has its own retention time tolerance, the data are searched in the same way as in any other experiment.

### Human and yeast benchmarking

#### Label-free data

LF no-plex human and yeast benchmarking data was searched with JMod (v2.0.0 timePlex beta) using the following commands: *{*–iso*}*, *{*–num iso 2*}*, *{*–lib frac 0.2*}*, *{*– no ms1 req*}*, *{*–atleast m 2*}*. LF 3-timePlex data was searched using the following commands: *{*–iso*}*, *{*–num iso 2*}*, *{*–lib frac 0.2*}*, *{*–timeplex*}*, *{*–num timeplex 3*}*, *{*–no ms1 req*}*, *{*– atleast m 2*}*. Experiments for evaluating timePlex carryover utilized the following commands:*{*– user rt tol*}* and *{*–rt tol 2*}* to set per-channel RT tolerances to 2 minutes rather than allowing dynamic settings.

#### plexDIA data

The 3-plexDIA human and yeast benchmarking data was searched with JMod (v2.0.0 timePlex beta) using the following commands: *{*–iso*}*, *{*–num iso 2*}*, *{*–lib frac 0.2*}*, *{*– no ms1 req*}*, *{*–atleast m 2*}*, *{*–plexDIA*}*, *{*–tag mTRAQ*}*. 9-plexDIA data was searched with JMod (v2.0.0 timePlex beta) with the following commands: *{*–iso*}*, *{*–num iso 2*}*, *{*–lib frac 0.2*}*, *{*–no ms1 req*}*, *{*–atleast m 2*}*, *{*–plexDIA*}*, *{*–tag PSMtag 9plex*}*.

#### Combinatorially multiplexed data

Combinatorial 3-timePlex and 3-plexDIA human and yeast benchmarking data was searched with JMod (v2.0.0 timePlex beta) using the following commands: *{*–iso*}*, *{*–num iso 2*}*, *{*–lib frac 0.2*}*, *{*–timeplex*}*, *{*–num timeplex 3*}*, *{*–no ms1 req*}*, *{*–atleast m 2*}*, *{*–plexDIA*}*, *{*–tag mTRAQ*}*. Combinatorial 3-timePlex and 9-plexDIA data was searched with JMod (v2.0.0 timePlex beta) with the following commands: *{*–iso*}*, *{*–num iso 2*}*, *{*–lib frac 0.2*}*, *{*–timeplex*}*, *{*–num timeplex 3*}*, *{*–no ms1 req*}*, *{*–atleast m 2*}*, *{*–plexDIA*}*, *{*–tag PSMtag 9plex*}*.

All non-multiplexed data and FDR benchmarking data was filtered at 1% precursor and protein FDR (Qvalue *<* 0.01 & Protein Qvalue *<* 0.01). The following filters were applied to all multiplexed data: Best tP specific channelQvalue *<* 0.01 & Qvalue *<* 0.5 & Protein Qvalue *<* 0.01. This allows for including precursors of less-confident assignment if a given timePlex channel has *≥*1 mass channel with Qvalue *<* 0.01. This effectively transfers confidence of precursor assignments across plexDIA channels within a timePlex channel, but not between timePlex channels.

### Searching the single-cell spectra

Single cell combinatorial 9×3 plexDIA & timePlex data was searched with JMod using an empirical spectral library generated from 30-cell equivalents of digested peptides from Raji, K562, and HEK293 cells. The following commands were used: *{*–iso*}*, *{*–num iso 1*}*, *{*–lib frac 0.2*}*, *{*– timeplex*}*, *{*–num timeplex 3*}*, *{*–no ms1 req*}*, *{*–atleast m 1*}*, *{*–plexDIA*}*, *{*–tag PSMtag 9plex*}*, *{*–use emp rt*}*.

### Sample preparation

#### Creating a mixed species standard for benchmarking

Benchmarking samples consisted of commercial Yeast (V7461, Promega) and Human K562 (V6951, Promega) protein digests. The dried digests were resuspended in 100 mM TEAB, pooled using their appropriate ratios to create Samples A, B, and C, each sample consisted of 25 *µ*g of total human material for each sample. The samples were split in two, one to remain label-free and the other to be chemically labeled with mTRAQ. The label-free samples were dried, and the other set of samples were labeled with mTRAQ Δ0 (SCIEX 4440015), Δ4 (SCIEX 4427698), and Δ8 (SCIEX 4427700) at a 2:1 ratio of 500 ng/*µ*L sample in 100 mM TEAB : stock concentration of tag. After addition of the label, the samples were left at room temperature to react for 2 hours. Hydroxylamine was added to a final concentration of 0.2% to quench the reaction at room temperature for 30 minutes. The samples were then dried with a speedvacuum, and then resuspended in 0.015% DDM and 0.1% formic acid to their appropriate concentrations. PSMtag benchmarks utilized for the 500 SPD evaluations were prepared and reconstituted as described by Specht *et al*^32^. PSMtag benchmarks used for dual-timePlex evaluations were prepared identically, except that the samples were three unique ratios of mixed-species peptide digests identical to the aforementioned Samples A, B, and C. Samples were labeled with the following PSMtag mass tags and pooled into plexDIA sets of 100 ng per channel: A (Δ0), B (Δ2), C (Δ4), B (Δ6), C (Δ8), A (Δ10), C (Δ12), A (Δ14), B (Δ16).

#### Cell culture

K562 (CCL-243, ATCC) and Raji (CCL-86 ATCC) cells were cultured in RPMI-1640 Medium (11875-080, Gibco). HEK293 (CRL-1573, ATCC) were cultured in DMEM (30-2002, ATCC). Cell culture media were supplemented with 10% fetal bovine serum (10439016, Gibco) and 1% penicillin-streptomycin (15140122, Gibco). Cells were grown at 37*^◦^*C and 5% CO_2_, and harvested within 15 passages.

#### 10x bulk Raji, K562, and HEK293 cells

Raji, K562, and HEK293 human cells were harvested from culture, washed 2x in PBS, and resuspended in water to a concentration of approximately 2,000 cells/*µ*L. The cell suspensions were subjected to mPOP^71^. In brief, the cell suspensions were frozen at -80°C, then heated at 90°C in a thermal-cycler for 10 minutes. TEAB was then added to the cell lysate to a concentration of 100 mM, and Trypsin Platinum (VA9000, Promega) and LysC Endoproteinase (90307, Thermo Fisher) were added to a concentration of 10 ng/*µ*L, and digested at 37°C for 18 hrs, speedvacuum dried. Peptides were resuspended at 1 *µ*g/*µ*L in a mixture of 436.6 mM DIPEA in DMSO (*∼*30 eq). 1 *µ*L of PSMtag label (6 *µ*g/*µ*L in DMSO) was added to 1 *µ*L (1 *µ*g) of peptides and incubated for 2 hours at room temperature then quenched with 1 *µ*L of 5% hydroxylamine. The labeled samples were resuspended in 24% ACN + 2% formic acid + 0.05% DDM in LC-MS water, and mixed to final concentrations of 10-cells per channel.

#### Single-cell sample preparation using nPOP

Single cells were prepared for proteomic analysis by nPOP as previously described by Leduc *et al.*^56,57^. In short, single-cells (in a volume of 0.3 nL) were sorted by CellenOne into 3 nL droplets of 100% DMSO on a fluorocarbon-coated glass slide to lyse. The lysed cells were digested with a mixture containing Trypsin Platinum (VA9000, Promega) at a concentration of 100 ng/*µ*L and LysC (90307, Thermo Fisher) at a concentration of 40 ng/*µ*L and 0.015% DDM (w/v) and 20 mM HEPES (pH 8.5), 5 nL of digestion mixture was dispensed onto each droplet. Peptides were then chemically labeled with PSMtag Δ0, Δ2, Δ4, Δ6, Δ8, Δ10, Δ12, Δ14 or Δ16 (6 nL/droplet) and subsequent addition of 200 mM TEAB buffer (6 nL/ droplets, pH 8.5) was added for a 2 hr incubation period. The labeling reaction was quenched with 5% Hydroxylamine (6 nL/ droplets). Single cells were picked up by the CellenOne, using 70% Acetonitrile in LC-MS water, into 9-plex sets, and dispensed into wells of a 384 well-plate. The pooled samples in the 384 well-plate were dried in a speed-vacuum then stored at -80°C until use, at which point they were resuspended in 24% ACN + 2% formic acid + 0.05% DDM in LC-MS water.

### Computational data analysis

#### Quantification

To stabilize and improve quantification across runs, methods, and within non-multiplexed and timePlex multiplexed data acquisitions, we developed a RT-dependent normalization in JMod that flattens the local median precursor intensities across retention time to each channel’s own median, separately for MS1 and MS2 quantities. Additionally for timePlex, this strategy regresses out the RT-dependent deviations across timePlex channels. plexDIA channels share one correction, leaving within-set ratios unchanged. JMod outputs both unnormalized and normalized (suffix ‘ rtnorm’) quantities.

All benchmarking experiments, multiplexed and non-multiplexed, were quantified at precursor-level using MS1 ‘plex Area rtnorm’ output from JMod, with the exception of mixed-species bulk 9-plexDIA PSMtag data which were quantified at MS2-level utilizing a function that requantifies each channel from fragments that are channel-unique, for example excluding super-positions of y-ions from peptides that are not C-terminally mass-tagged. The function additionally accounts for carryover of isotopic envelopes across 2-Da offset channels. This MS2-level quantification (‘co-eff ChannelUnique CommonApex rtnorm’ and ‘coeff ChannelUnique Area rtnorm’) is output by JMod.

All quantitative benchmarking was performed using the median value across triplicate samples. For example, in the case of LF no-plex 9 LC-MS runs were required to analyze triplicates of Samples A, B, and C, respectively; 3 runs are required to analyze triplicates of the same samples for 3-plexDIA or 3-timePlex; and finally, a single run was used to analyze triplicates of the same samples for 3×3 plexDIA & timePlex. This way, medians were computed from the same number of samples regardless of the method. This was performed for all quantitative comparisons unless otherwise specified.

#### Entrapment to assess false discoveries of timePlex

An empirical library generated by DIA-NN^15^ for human and yeast LF peptides was created, as well as a predicted library for *A. thaliana*. Precursors from the *A. thaliana* predicted library were randomly sampled to have the same total number as in the empirical human and yeast library. The two equivalently sized libraries were merged, and this merged library was used to search LF 3-timePlex data which had missing yeast in one time channel, and *A. thaliana* missing from all time channels. Data were analyzed at 1% FDR.

#### Single-cell protein quantification, imputation, batch correction, and PCA

Negative controls were prepared and acquired alongside real single-cells. The distribution of summed MS1 signal normalized to the median of each timePlex channel of each run for negative controls was used to separate (un)satisfactorily prepared single-cells. The cutoff was selected as +2 SD above the negative-control mean. Of remaining single-cells, precursors were quantified at MS1 level using ‘plex Area rtnorm’. These precursor values were median normalized to account for absolute differences in intensities, then each precursor was normalized to the mean of its respective plexDIA set, and transformed back to absolute intensities by multiplying by the mean of absolute intensity of each precursor. MaxLFQ quantitation was then applied on these normalized values using the DIANN R package^15,72^. Protein quantities were normalized by their median abundances for each cell to account for absolute differences, then normalized across cells by the mean. Protein data were batch-corrected using ComBat^73,74^ to account for PSMtag label and LC column biases. These data were then filtered to the subset of proteins quantified in *≥*50% of single-cells and intersected with bulk 10x data to compute a Pearson correlation for assessing quantitative agreement of averaged single-cells and bulk. The log_2_-transformed protein data was imputed using a weighted kNN approach (k=10), where the weights were scaled based on euclidean distances to its nearest neighbors. And finally, principal components were computed from imputed single cell and 10x bulk protein data where the variance of each protein was scaled by the sum of squared protein-protein correlations.

## Data and code availability

All data were reported and deposited in accordance with the community guidelines^75^. Raw data, spectral libraries, search outputs, instrument methods, and meta data are available at MassIVE: MSV000103762. Code for repeating downstream analyses is available at github.com/ParallelSquared/timePlex.

## Acknowledgments

We thank Andrew Leduc for work and suggestions that improved sample preparation by nPOP for PSMtag-labeled single-cells. PTI is a Convergent Research Focused Research Organization (FRO) and has received support from Eric and Wendy Schmidt as well as Griffin Catalyst. Initial conceptualization occurred when J.D. and N.S. were supported by a MIRA award from the NIGMS of the NIH (R35GM148218) to N.S.

## Author contributions

**Experimental design**: J.D., H.S., and N.S.

**Data analysis**: J.D., K.M.D., D.J.G., N.W. and N.S.

**LC-MS/MS**: J.D., S.N.S.

**timePlex-enabling design**: J.D., H.S., J.J.S., L.C., and X.X.

**Sample preparation**: J.D., L.W., P.S., M.P.A., and M.Y.

**Raising funding**: N.S. and H.S.

**Initial draft**: J.D., and N.S.

**Writing**: All authors approved the final manuscript.

## Extended Data Figures

**Extended Data Fig. 1.**
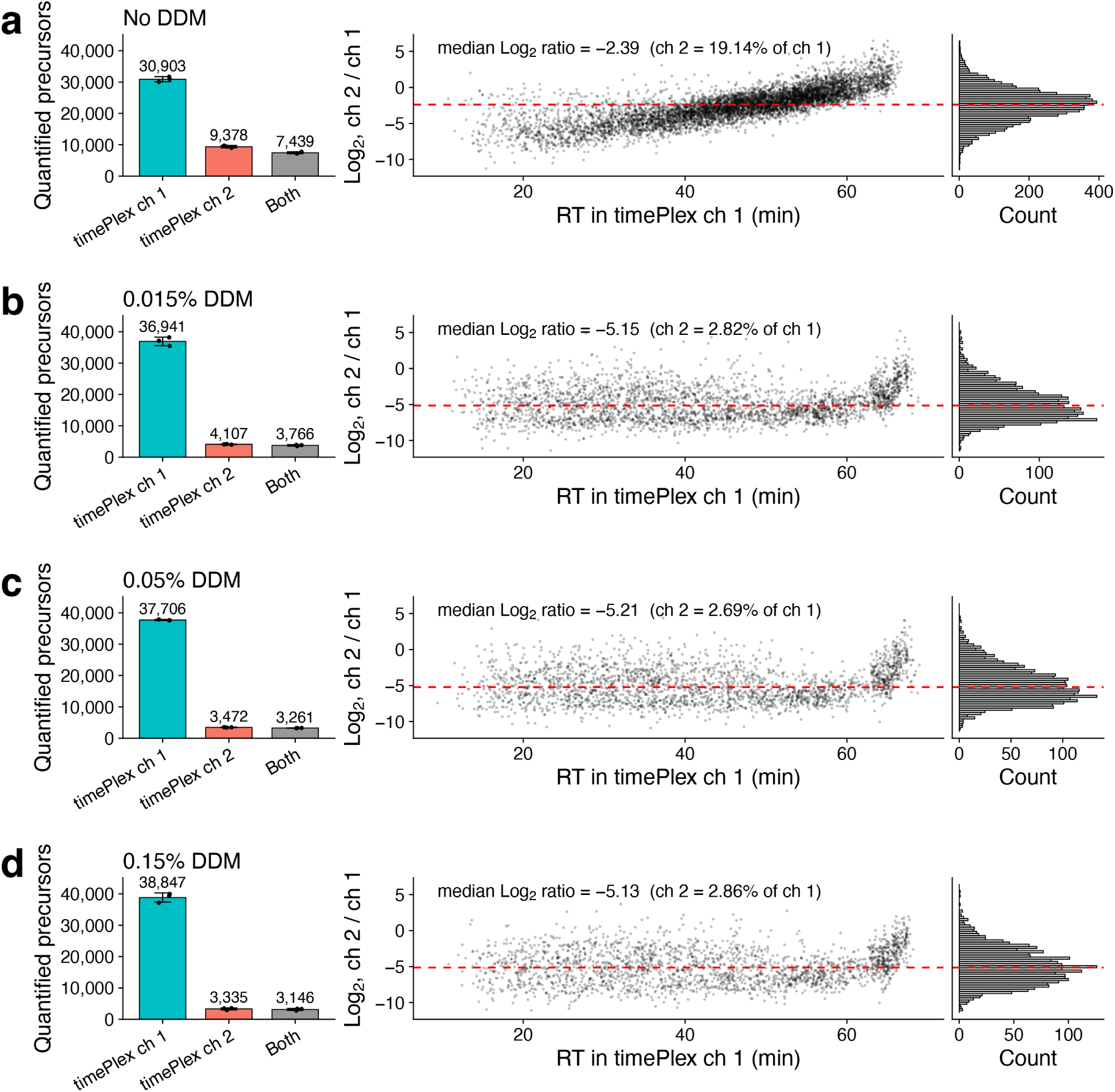
Assessing carryover between timePlex channels. The timePlex sample loading performed in this work occurred through a shared sample-loop and transfer lines. In some cases, this may result in unspecific binding of peptides onto these surfaces which will flow unintended into the third column during the active gradient. To mitigate this, samples can be resuspended in a mass-spec compatible detergent (such as DDM) to reduce carryover. To assess the carryover, LF non-multiplexed data were searched as a 2-timePlex, attempting to quantify carryover into the unintended time channel. **a** 10 ng samples which were not resuspended in DDM bound to common flow path and carried over to an unintended time-channel (“ch 2”). This resulted in the unintended channel quantifying approximately 30% of the number of precursors as the intended time-channel, of which the precursors had a median of 19% the intensity, with increasing intensity at higher hydrophobicity – consistent with unspecific binding to the common flow path. **b** The same experiments were performed with 0.015% DDM, **c** 0.05% DDM, and **d** 0.15% DDM, all of which produced approximately 10% of the number of quantified precursors in the unintended channel. Of these precursors, their intensities were *<* 3% of the intended time-channel, suggesting DDM of 0.015% concentration is sufficient to bring column carryover to a low level.

**Extended Data Fig. 2.**
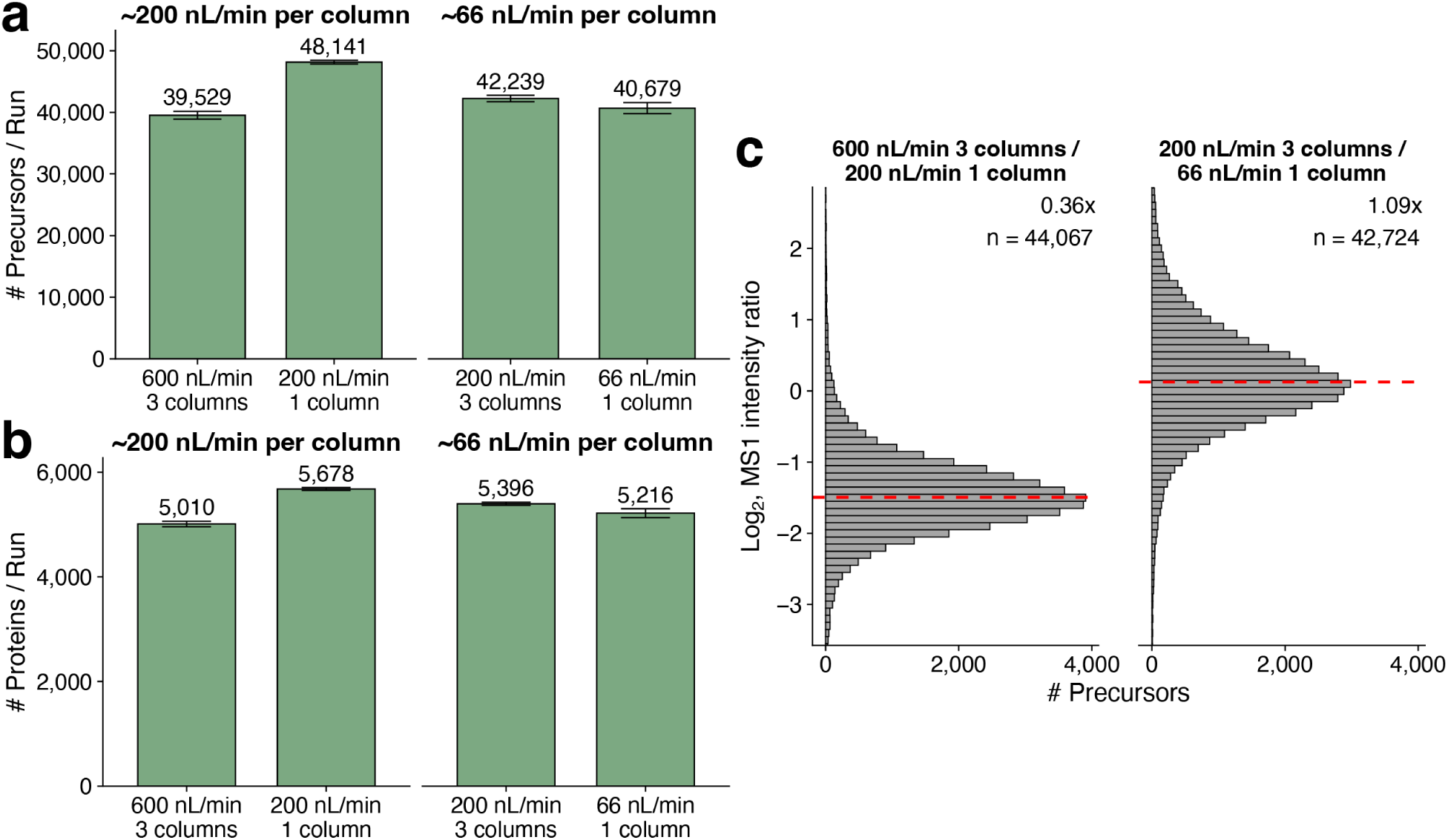
Investigating the effect of the 3 column separate emitter approach (background spray). Here we investigate how the 3-column setup alone (not the timePlex data-type) impacts ion intensities and coverage relative to standard proteomics acquisition from a single emitter. **a** Bar plots show the number of precursors identified for 200 nL/min flow rate per column, achieved by flowing through a single column or through 3 columns splitting 600 nL/min. In addition 66 nL/min/column achieved through a single column or 200 nL/min split over 3 columns. Error bars represent ± standard error (*n* = 3 runs). **b** Number of proteins identified per run. **c** Ratio of precursor MS1 intensities when 3 columns are spraying a total of 600 nL/min (however, only 1 column is actively separating a sample), or 1 column is spraying 200 nL/min. The 200 nL/min/column 3-column setup results in approximately 3-fold lower MS1 intensities. However, at the lower flow rate of 66 nL/min/column, the 3-column approach was similar intensity to the single-column approach. The red dashed line indicates the median of the distribution.

**Extended Data Fig. 3.**
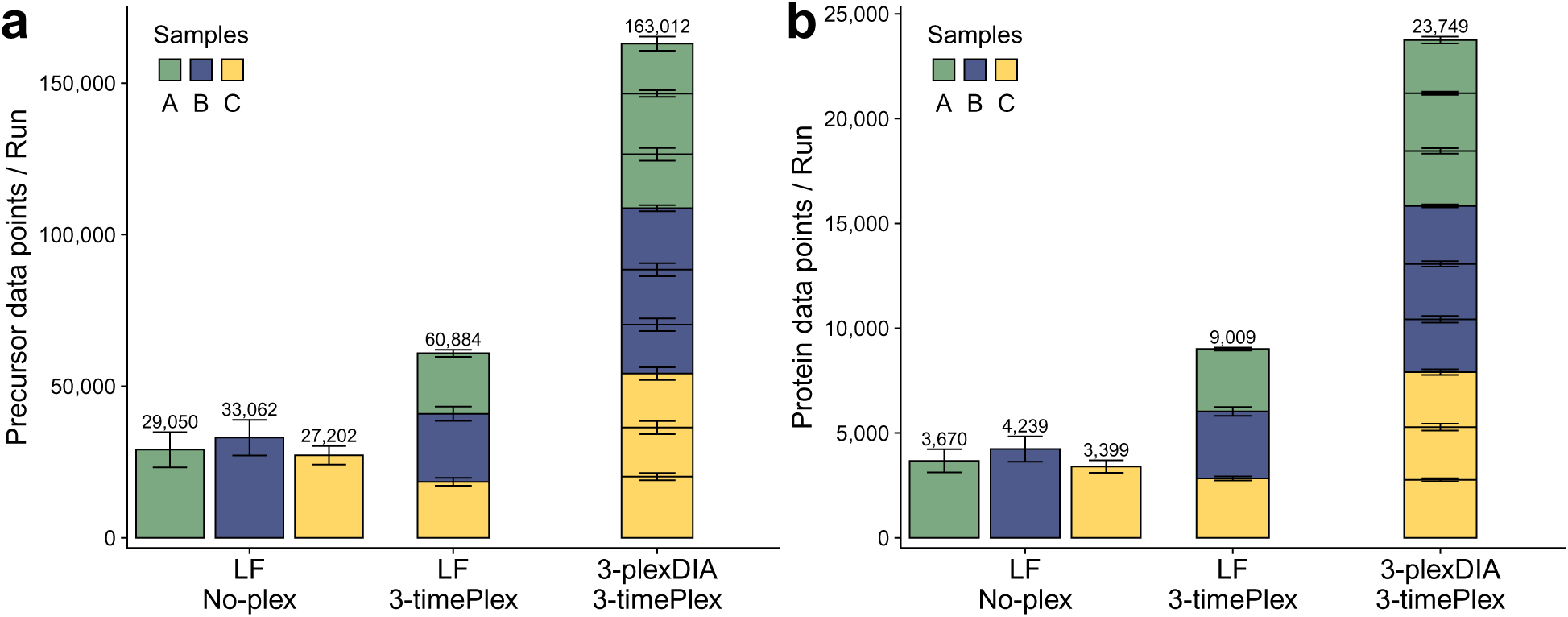
Precursor and protein coverage for timePlex and combinatorial plexDIA & timePlex using merged flow approach. All data were acquired with the same experimental setup whereby columns (Thermo *µ*PAC Neo 50 cm columns) were joined through a union to spray from a single emitter, including LF no-plex, for purposes of controlling for the experimental implementation and evaluating the timePlex datatype. a-b Number of precursor data points a and protein data points b identified per run for each data type; error bars represent ± standard error.

**Extended Data Fig. 4.**
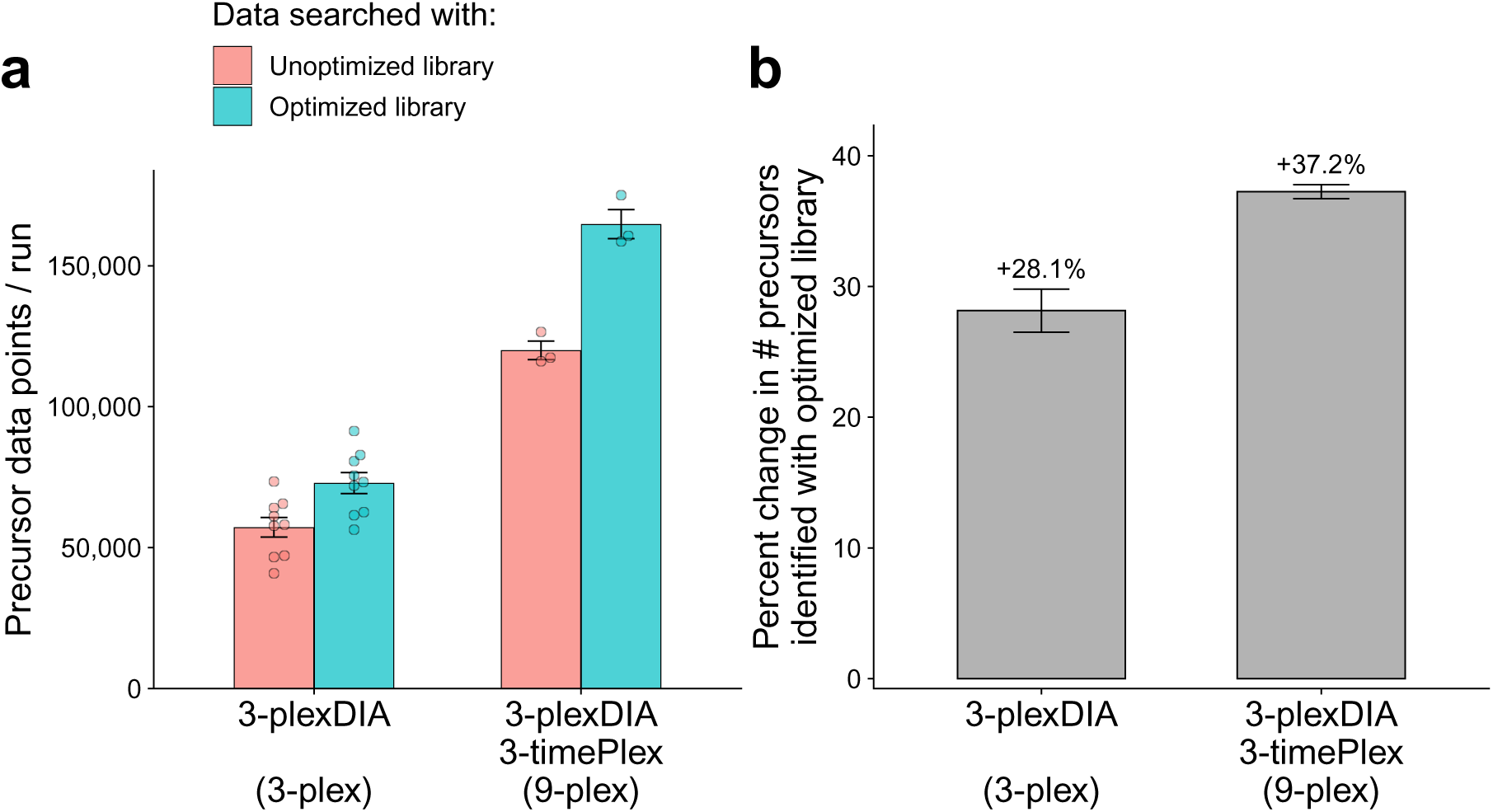
Spectral library quality becomes increasingly important at high-plex. **a** Triplicates of each plexDIA or combinatorial plexDIA & timePlex set were acquired, resulting in 9 runs for 3-plexDIA and 3 runs for 3×3 plexDIA & timePlex. Shown is the number of precursors identified for each run, searched with an unoptimized library (red) or mTRAQ optimized library (blue). To enable a fair comparison of the libraries, the two libraries were intersected which resulted in smaller library sizes overall and thus reduced total coverage. **b** The same data, but shown as the change in the number of precursors identified as a result of searching data with an unoptimized library or mTRAQ optimized library. Searching data with an optimized spectral library results in a greater percentage increase of identifications at higher plex.

**Extended Data Fig. 5.**
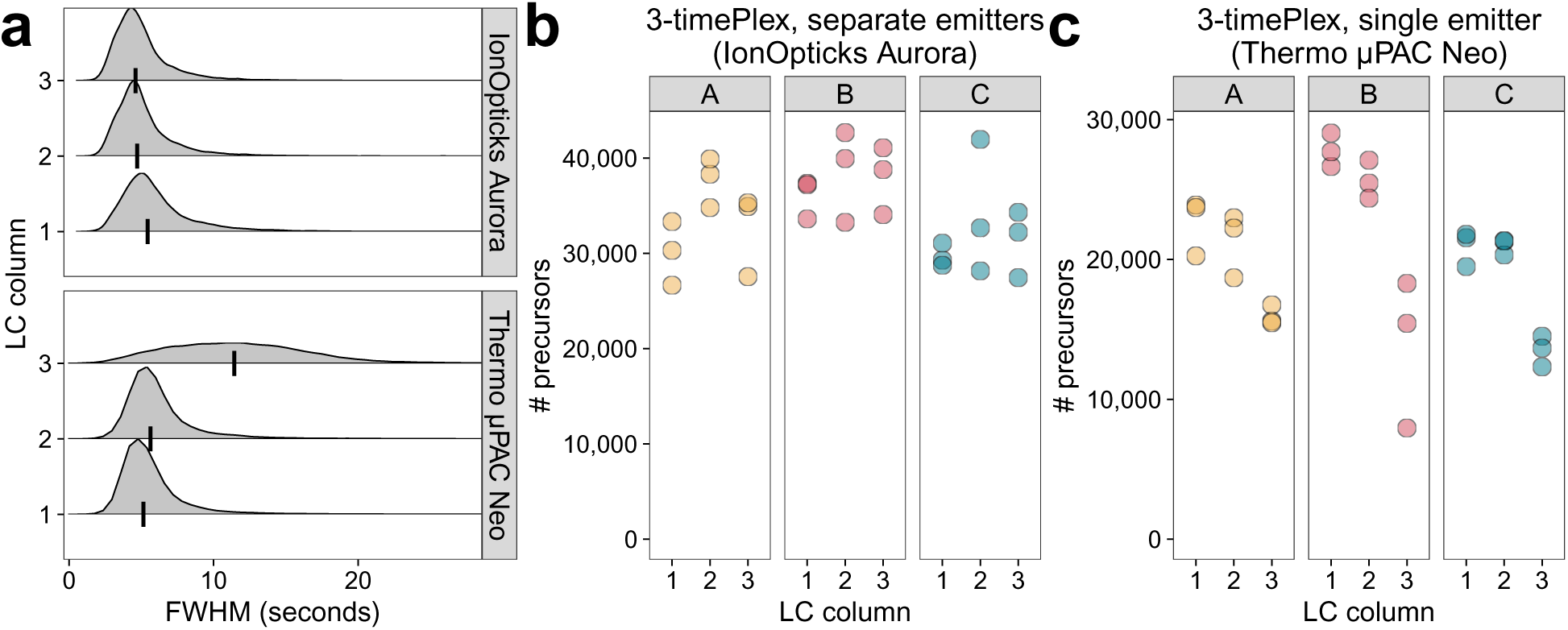
Investigating column-specific differences with timePlex. **a** Full width half max (FWHM) of precursor elutions was intersected across columns and implementations for the ‘separate emitters’ approach using IonOpticks Aurora 25cm x 75*µ*m columns, or the ‘single emitter’ approach using Thermo *µ*PAC Neo 50 cm columns, and plotted as density plots; a vertical black line marks the median of the distributions. **b** The ‘separate emitters’ approach using IonOpticks columns produced similar precursor coverage across columns. In the figure, each point represents a replicate and each facet corresponds to a sample (A, B, C). **c** Similar trends are observed for the ‘single-emitter joined flow’ approach using Thermo *µ*PAC Neo columns, however the 3rd column consistently under-performed the first two.

**Extended Data Fig. 6.**
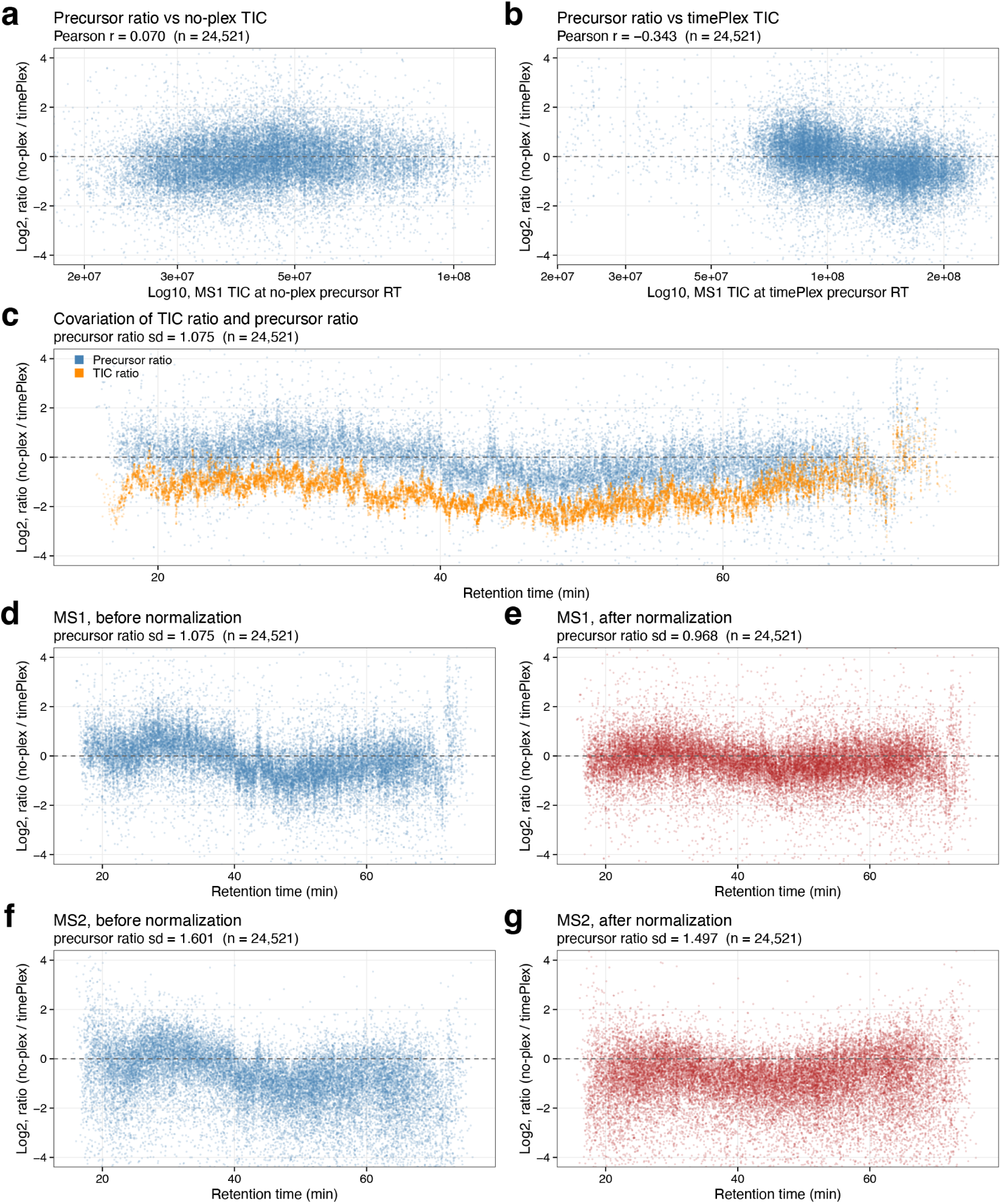
Investigating factors influencing timePlex and no-plex quantitative agreement and corrections. **a** Log_2_ ratio of a LF no-plex run and a LF 3-timePlex run vs the log_10_ TIC at the RT when the no-plex precursor eluted, r=0.070. **b** Same as previous panel, but TIC corresponds to the timePlex run, r=-0.343. **c** Plotting the precursor ratios and TIC ratios vs RT; text indicates the standard deviation (sd) of the precursor ratios. **d–g** Precursor ratios quantified at MS1 (**d, e**) and MS2 (**f, g**), shown before (**d, f**) and after (**e, g**) the normalization strategy introduced in this work.

**Extended Data Fig. 7.**
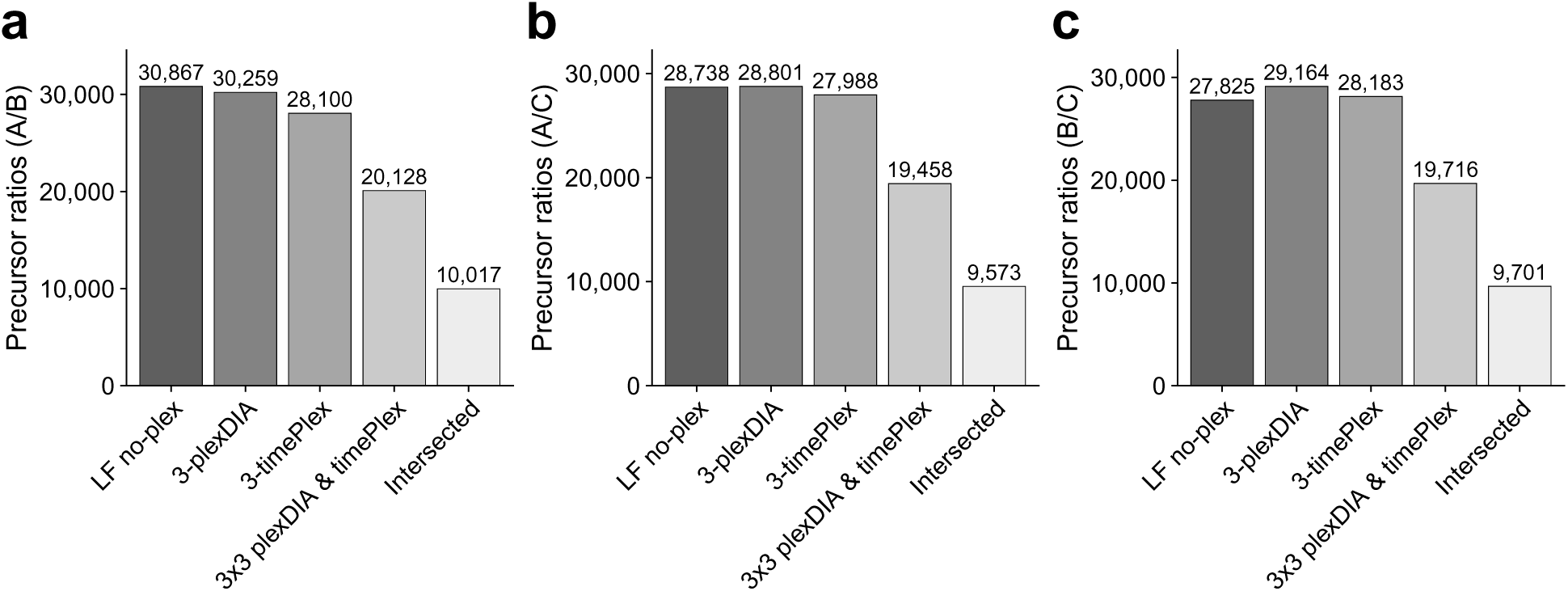
Number of precursor ratios quantified across methods. Number of precursor ratios quantified between samples A and B (**a**), A and C (**b**), and B and C (**c**) for label-free (LF no-plex), 3-plexDIA, 3-timePlex, and 3×3 plexDIA & timePlex, using the ‘separate emitters’ approach. A ratio was counted as quantified if the precursor was detected in both samples in at least 2/3 replicates. The intersect is also plotted for the number of precursor ratios that were quantified in common across all four methods.

**Extended Data Fig. 8.**
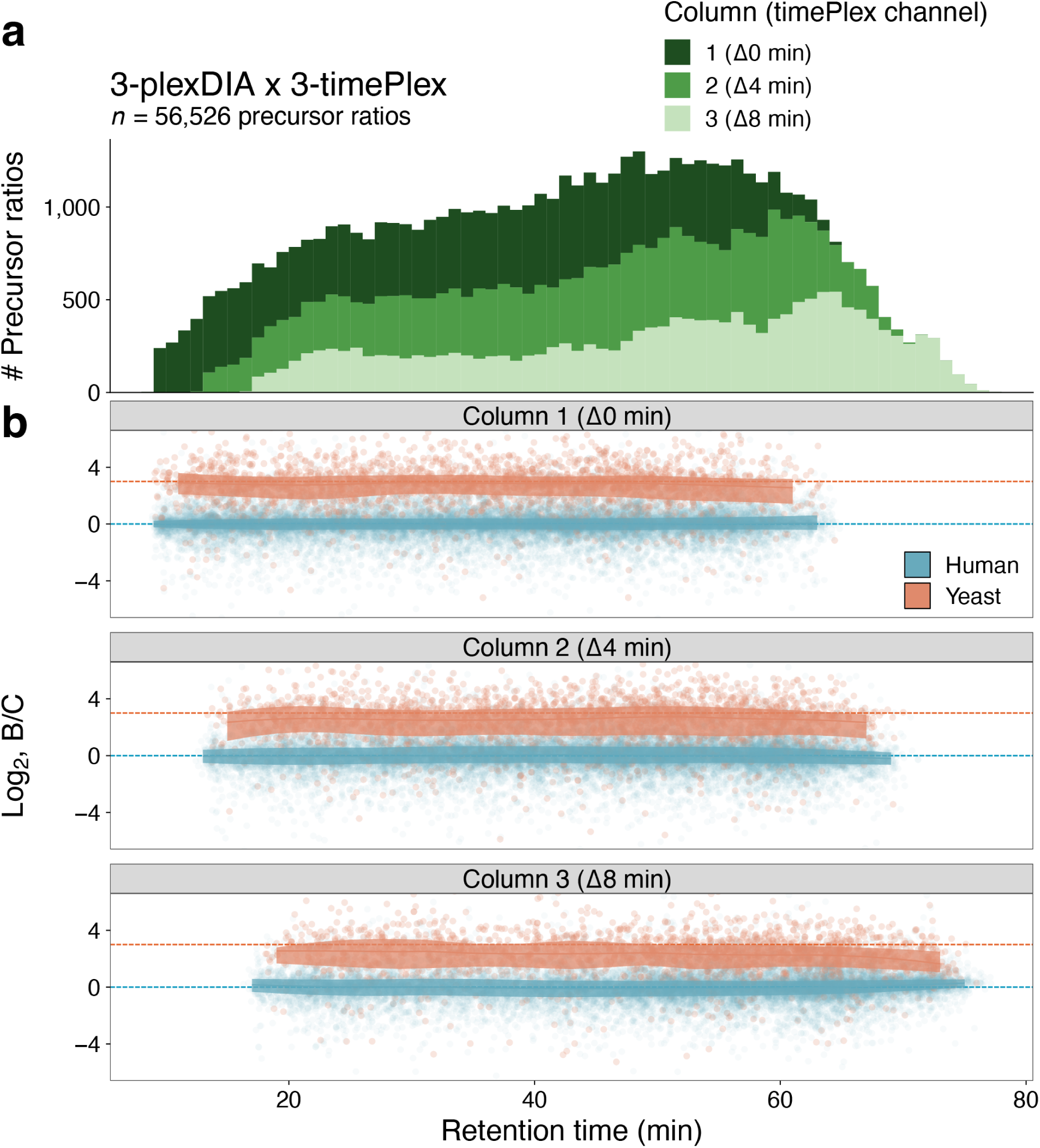
Evaluating quantitative accuracy across the gradient for 3×3 plexDIA & timePlex. **a** Number of precursor ratios quantified between samples B and C for a single 3×3 plexDIA & timePlex run, shown as a stacked histogram colored by timePlex channel. **b** Precursor ratios between samples B and C within each plexDIA set, plotted against retention time for each timePlex channel of the run. Orange and blue points represent yeast and human precursor ratios, respectively. Dashed lines indicate the expected fold changes between samples, and shaded ribbons denote interquartile ranges.

**Extended Data Fig. 9.**
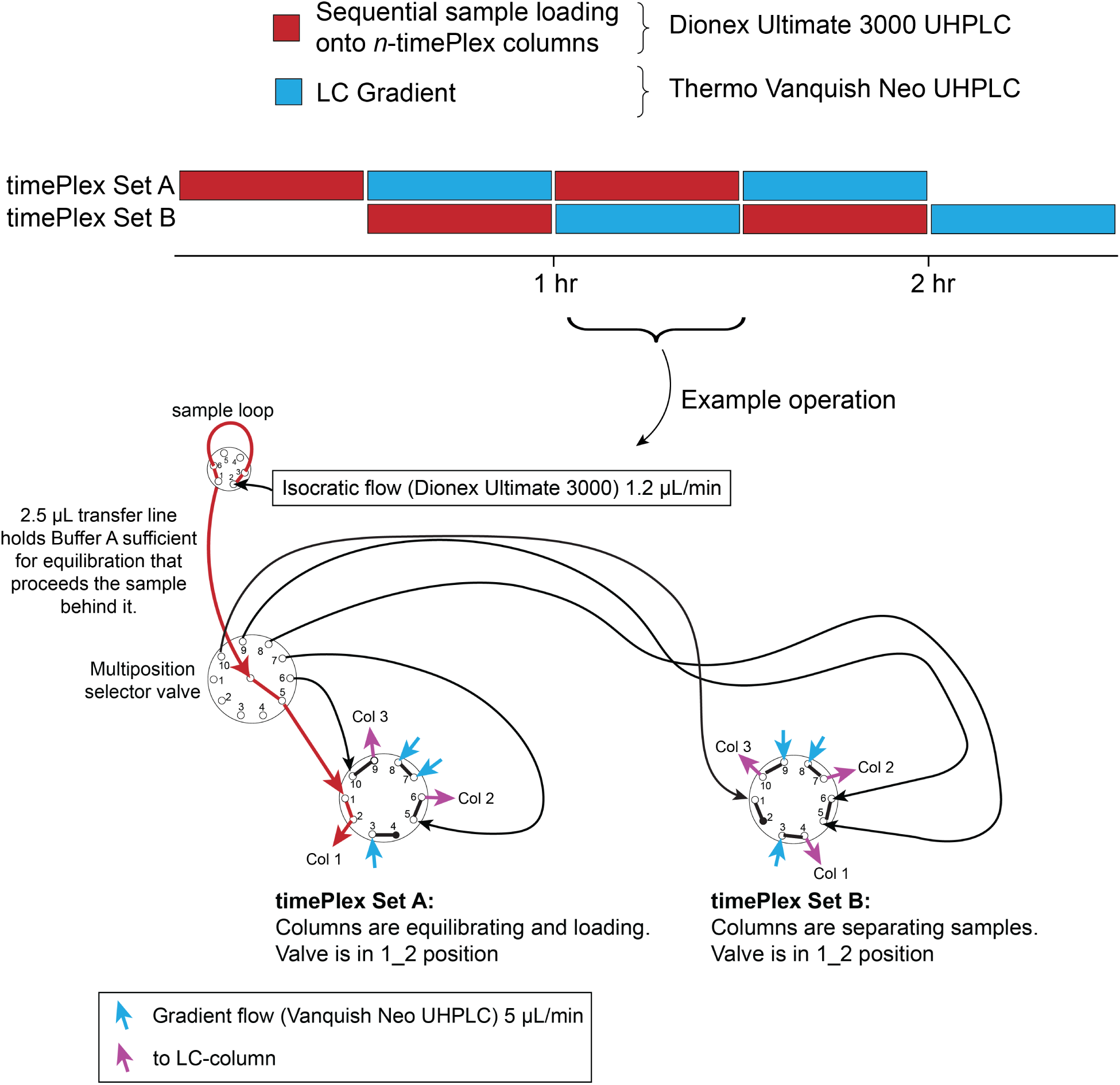
Dual-timePlex implementation. Dual-timeplex is implemented in this work through the use of two UHPLC systems, a Dionex UltiMate 3000 and a Thermo Vanquish Neo. The UltiMate 3000 performs column equilibration and sample loading of all columns, and simultaneously the Vanquish Neo delivers the gradient to the active timePlex set. All flow (sample loading and gradient delivery) is routed through two 10-port 2-position valves in the column compartment of the UltiMate 3000. The UltiMate 3000 delivers sample to each respective column as routed by a high pressure multiposition selector valve. The UltiMate 3000 relays are used to trigger the Vanquish Neo to start its gradient, begin MS acquisition, and switch voltage output through a high voltage relay. An additional 2-position valve (not shown) is used on the Thermo Vanquish Neo to deliver gradient flow to either Set A or Set B.

**Extended Data Fig. 10.**
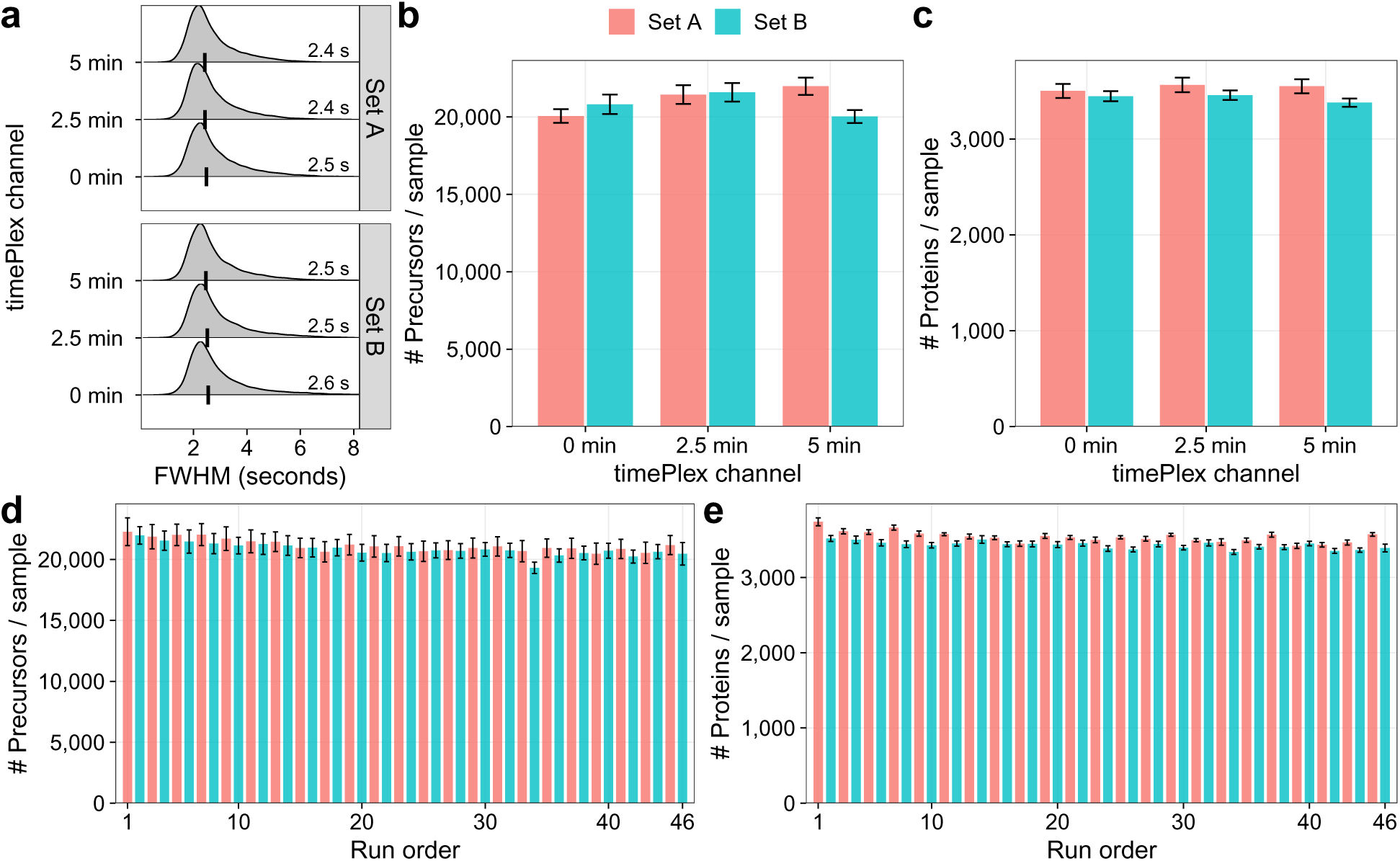
Evaluating consistency of combinatorial 9×3 plexDIA & dual-timePlex. **a** The distribution of intersected precursor full width at half max (FWHM) is shown in seconds for each of the 6 LC-columns used; the medians are marked with black tick marks and written in each respective panel. **b** Average number of precursors identified at 1% FDR for each timePlex channel and set. Red and Blue represent timePlex Set A and Set B, respectively. **c** Average number of proteins identified per sample at 1% FDR for each timePlex channel and timePlex set. **d-e** Average number of precursors **d** and proteins **e** identified per sample for each run over the 24 hour-span. All error bars represent standard deviation.

## Supplementary Figures

**Supplementary Fig. 1.**
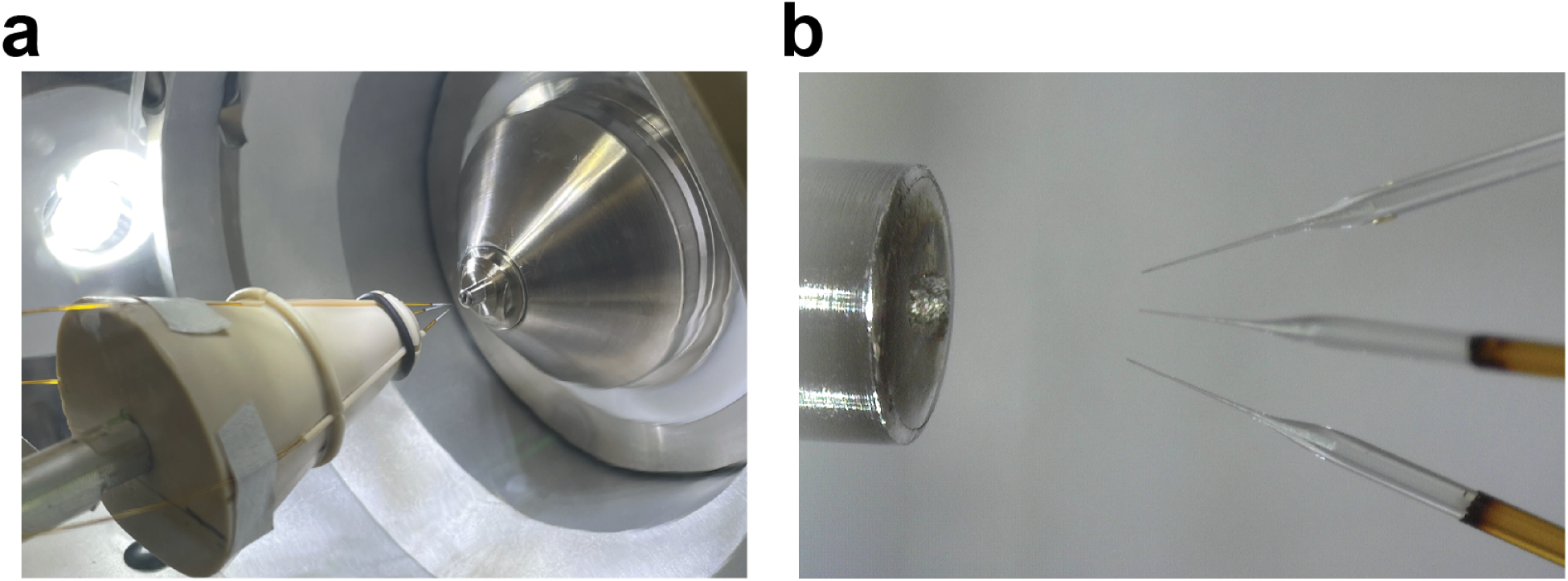
’Separate emitter’ setup for timePlex. **a** Photo of three 25 cm x 75 micron ID IonOpticks columns held in place by a PEEK cone, positioned at the mass spectrometer inlet. **b** Zoomed-in photo of the three emitters and the mass spectrometer inlet.

**Supplementary Fig. 2.**
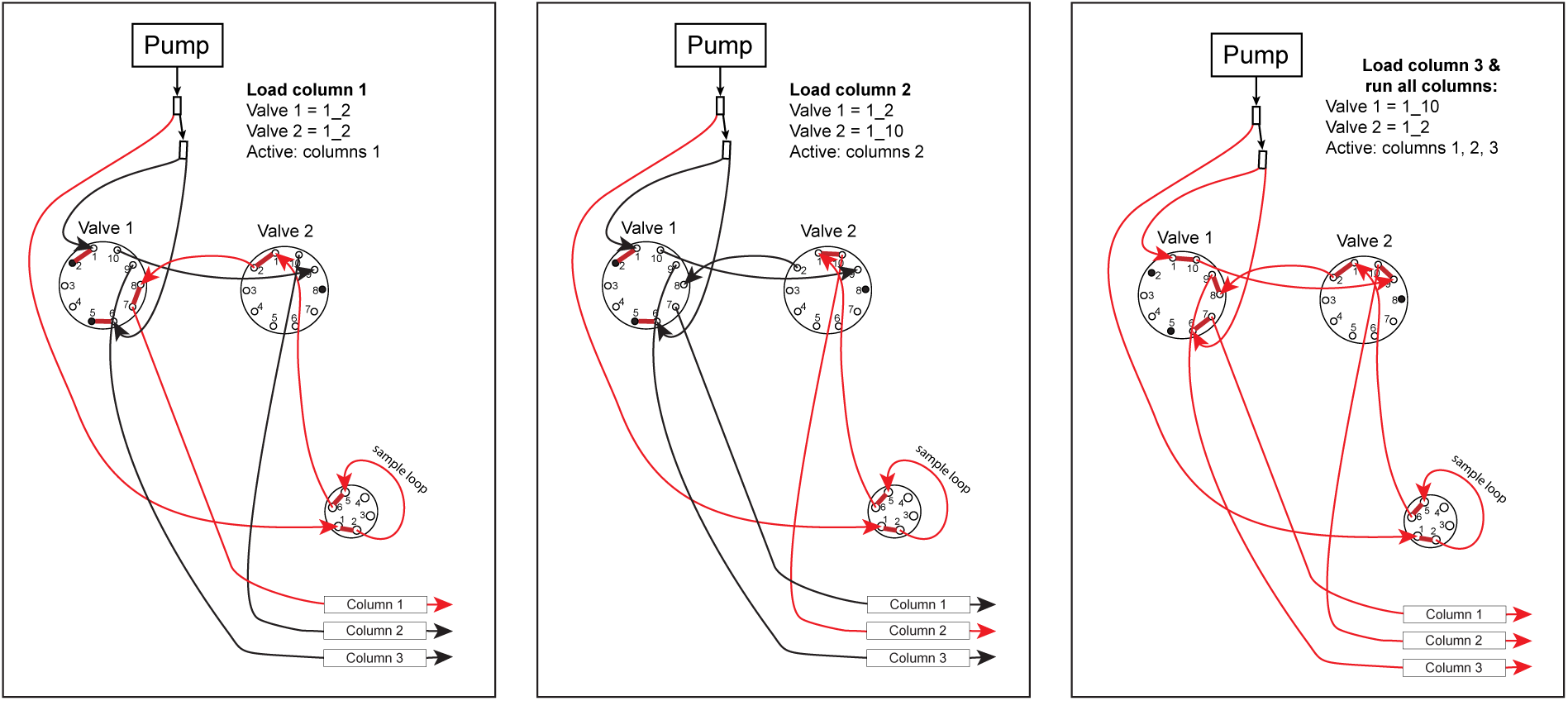
Implementing 3-timePlex on Dionex UltiMate 3000 HPLC. The diagram illustrates how samples are loaded sequentially for each column and then run together. Red indicates where net flow occurs. Flow is bifurcated from the NC pump, one line leads to the sample loop to push sample onto a given column, and the other line is bifurcated again to ports 1 and 6 of Valve 1. Plugs are utilized to maintain pressure to push sample through the loop during column loading. The valve-positions required to achieve this are noted on each panel of the diagram. Flow-rate of the methods are adjusted to account for the number of columns actively receiving flow. In this implementation, columns 1 and 2 are loaded at 300 nL/min to keep the backpressure of the system low as only a single column is active at this time; this produces approximately 1.5x the backpressure as three columns run at 600 nL/min, which occurs during the ‘Load column 3 and run all columns’ phase.

**Supplementary Fig. 3.**
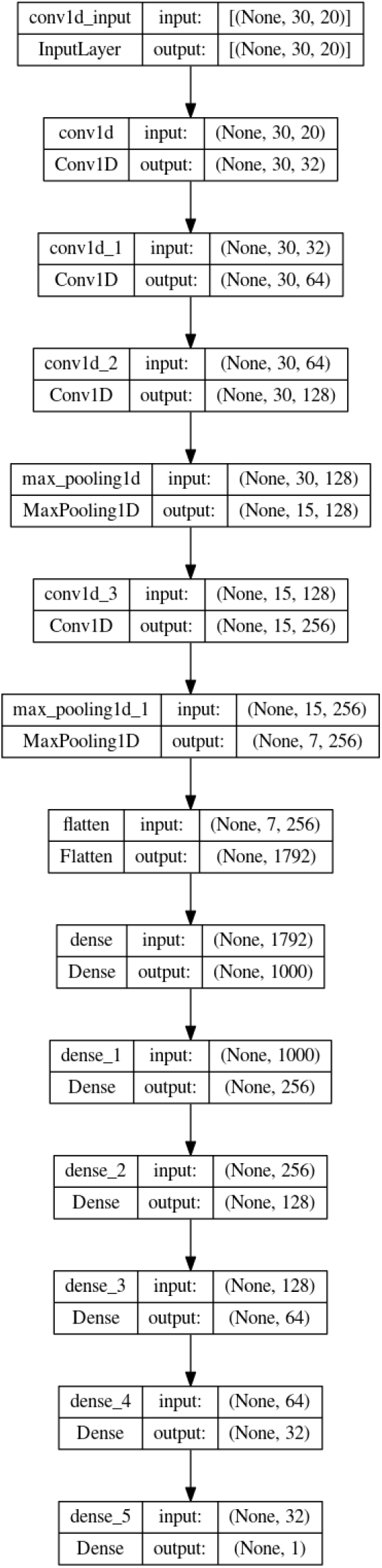
Model architecture for predicting retention times. The model is a hybrid CNN and fully connected feedforward DNN which takes inputs of shape (30, 20), 30 being the peptide length and 20 being placeholders for each amino acid. There are 4 1D convolutional layers with kernel size = 4. The number of filters applied at each convolutional layer was set to double, starting at 32 and ending with 256. Max pooling = 2 was used for the final two layers. The final convolutional layer was flattened and fed into a fully connected feed-forward deep neural network with 1000, 256, 128, 64, and 32 nodes at each layer. Swish activation functions were used at all layers, with the exception of the final layer which was linear for computing the final RT output.

**Supplementary Fig. 4.**
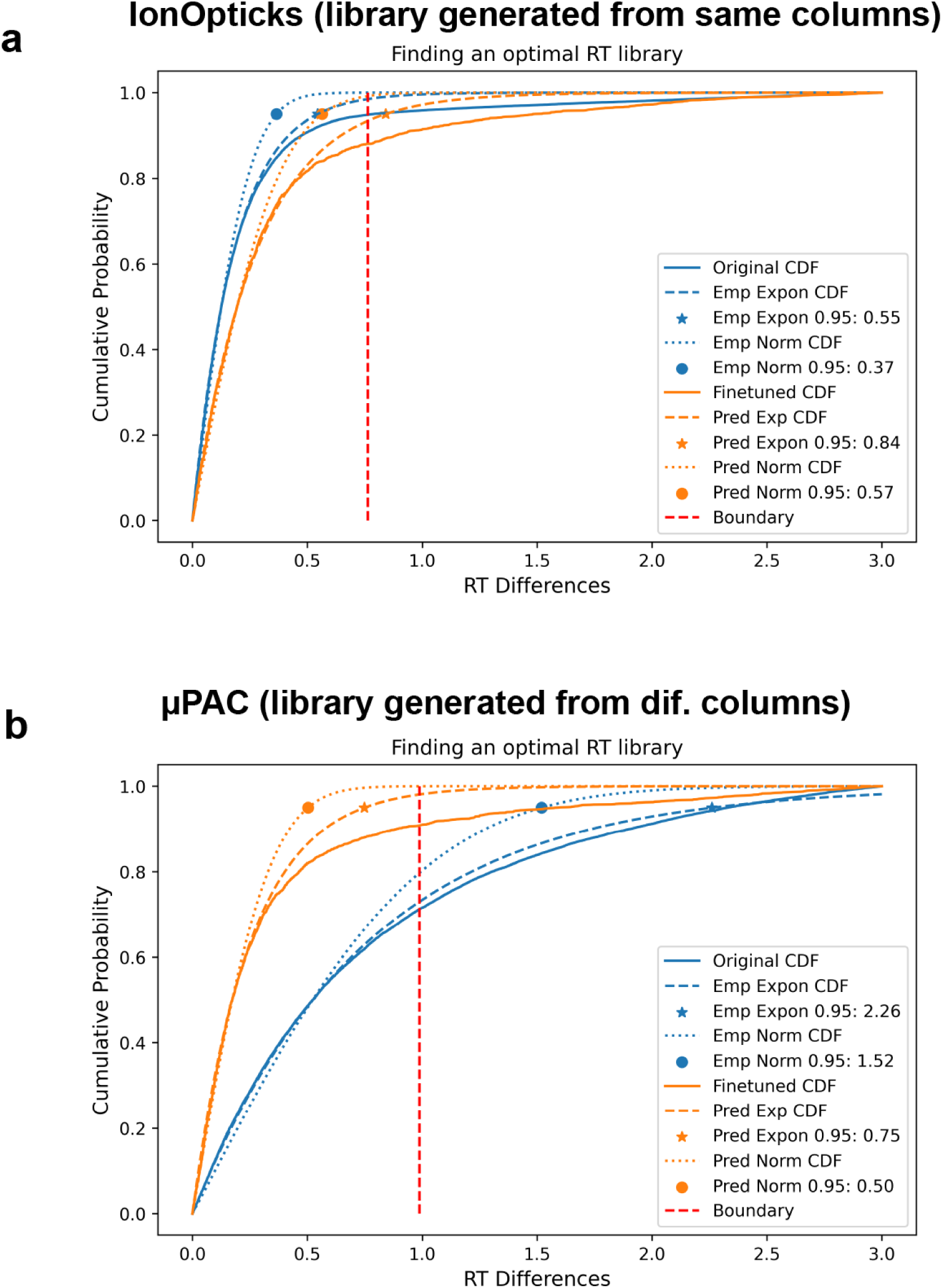
Setting retention time tolerances. JMod searches benefit from thresholding on a tightly set RT range, so a peptide is only considered within a specific RT window and not throughout the run. This is especially important in timePlex, where peptides must be correctly associated to their sample origin. To set appropriate RT windows, we compute a cumulative distribution on the validation dataset, comparing the original library RTs with the fine-tuned RTs. We fit the cumulative distributions as Gaussian or exponential, and choose the fit that best matches the actual distribution, setting the RT threshold at *±*4 SD. **a** Fine-tuning is not necessary for data where the library has accurate RTs, as is the case here where the libraries were acquired shortly after the benchmarking runs and originate from the same columns used in the benchmarking data. **b** However, in cases where the library RTs do not properly model the empirical RTs, fine-tuning (orange) is highly beneficial, allowing for a smaller range to consider a peptide eluting. Here the library originates from RTs from IonOpticks columns, leading to a large discrepancy between what is empirically observed on *µ*PAC columns (blue).

